# Geometry-based BERT: an experimentally validated deep learning model for molecular property prediction in drug discovery

**DOI:** 10.1101/2024.12.24.630211

**Authors:** Xiang Zhang, Chenliang Qian, Bochao Yang, Hongwei Jin, Song Wu, Jie Xia, Fan Yang, Liangren Zhang

**Author notes:** Correspondence should be addressed to J.X. or F.Y.; Dr. Jie Xia, Institute of Materia Medica, Chinese Academy of Medical Sciences & Peking Union Medical College, Beijing, China; Dr. Fan Yang, Department of Automation Xiamen University, Xiamen, China.

## Abstract

Various deep learning based methods have significantly impacted the realm of drug discovery. The development of deep learning methods for identifying novel structural types of active compounds has become an urgent challenge. In this paper, we introduce a self-supervised representation learning framework, i.e., GEO-BERT. GEO-BERT considers the information of atoms and chemical bonds in chemical structures as the input, and integrates the positional information of the three-dimensional conformation of the molecule for training. Specifically, GEO-BERT enhances its ability to characterize molecular structures by introducing three different positional relationships: atom-atom, bond-bond, and atom-bond. By benchmarking study, GEO-BERT has demonstrated optimal performance on multiple benchmarks. We also performed prospective study to validate the GEO-BERT model, with screening for DYRK1A inhibitors as a case. Two potent and novel DYRK1A inhibitors (IC_50_: <1 μM) were ultimately discovered at a hit rate of 10%. Taken together, we have developed the Geometry-based BERT model for molecular property prediction and proved its practical utility in early-stage drug discovery.

**Graphical Abstract:** 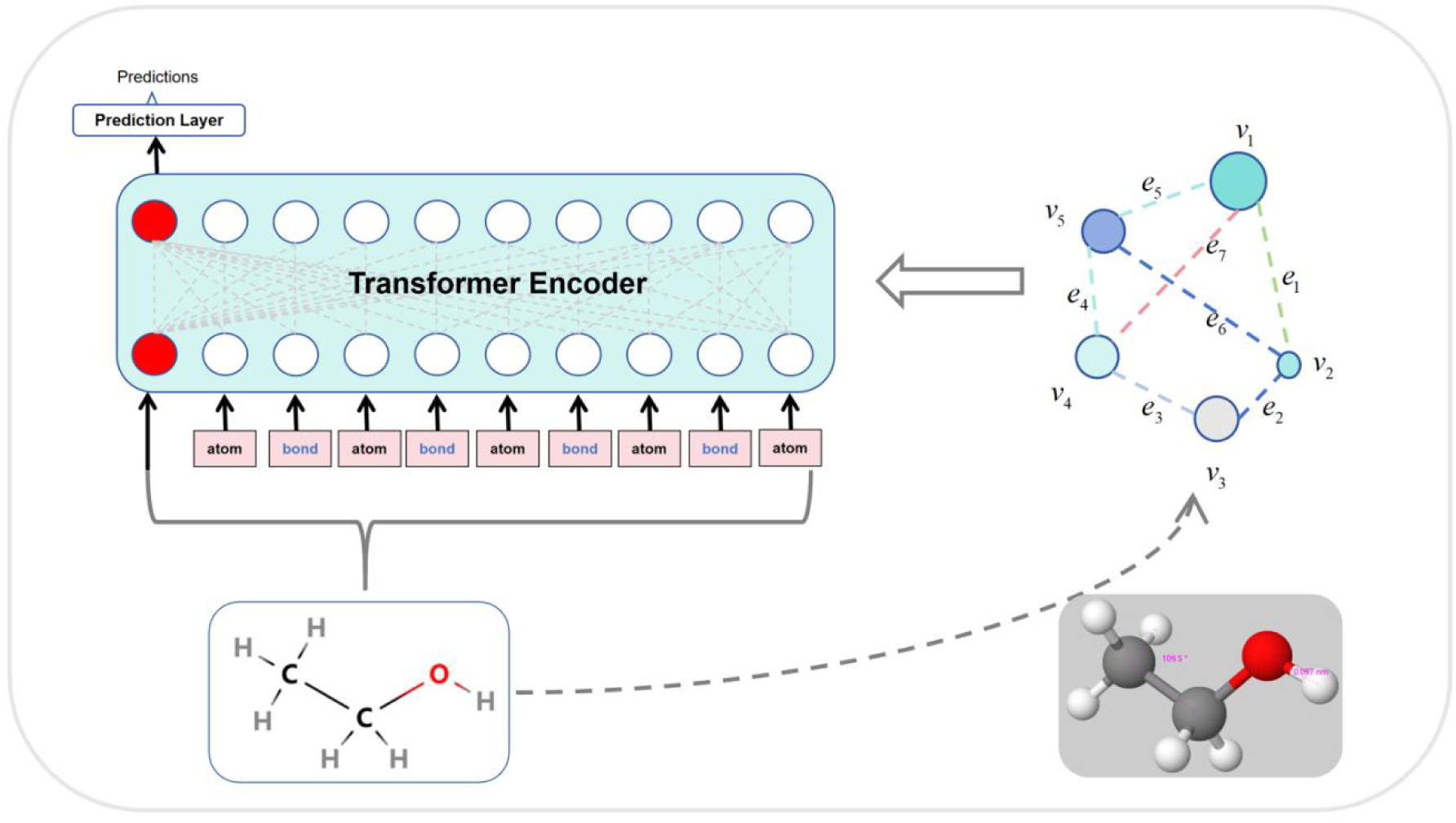

GEO-BERT is a model pretrained on large-scale drug molecule data, while improving the accuracy of property prediction by using three-dimensional structural information within the molecule.

## Introduction

In the history of drug discovery, pharmaceutical scientists often apply high-throughput screening to identify hit compounds^[1]^. However, it is well known that this technique is time-consuming and thus may significantly impede new drug discovery^[2]^. With technical advances, virtual screening has become an indispensable strategy to accelerate new drug research and development, as it is able to identify novel hits efficiently in a short period of time. As for virtual screening, normally a computational model is established and then used to filter a large-scale chemical library so as to select potentially active compounds from the library, and eventually discover truly active compounds through experiments^[3]^. As one of the aforementioned computational models, molecular property prediction based on machine learning has been widely used for virtual screening and drug discovery^[4]^.

Featurization of molecules is an essential step of machine learning-based molecular property prediction. It is conventionally achieved by the calculation of molecular fingerprints (e.g. ECFP^[5]^, MACCS^[6]^), or descriptors (e.g. structural descriptors, physicochemical property descriptors^[7]^). However, due to the fact that the featurization rules of molecular fingerprints and descriptors are based on human experience, these methods often have the following drawbacks: (1) Feature engineering requires manual design of molecular fingerprints/descriptors, relying on the knowledge and experience of domain experts^[8]^. (2) Different molecular fingerprints or descriptors in machine learning can bring about different performance, making it difficult to find the most suitable molecular fingerprints/descriptors for each machine learning algorithm^[9]^.

With the emergence of deep learning, new algorithms that do not require feature engineering have been developed, including Recurrent Neural Network (RNN), Convolutional Neural Network (CNN). These algorithms directly learn from end to end and train models based on the original molecular representations. It has been proved highly effective in handling diverse molecules^[10]^. According to the format of molecular representations, corresponding deep learning algorithms can be chosen and leveraged. For instance, the SMILES string representation of molecules are applicable as input of the algorithms such as Recurrent Neural Network (RNN) and Transformer^[11]^. The graph representations of molecules are suitable for Graph Neural Networks (GNN)^[12]^. GNN models usually employ message passing mechanism, enabling each node of the graph to aggregate information from its neighboring nodes. Each node can be additionally represented by the properties such as atomic type and charge. However, these supervised deep learning methods are faced with insufficient generalization, due to the limited data with labels^[13]^.

Natural language processing (NLP) provides an effective way to address the aforementioned issues. In recent years, Bidirectional Encoder Representations from Transformers^[14]^ (BERT) models attracted much attention in the field of molecular property prediction. By using its masking strategy and bidirectional attention, BERT models could achieve excellent generalization performance. BERT model building includes extensive pre-training with large-scale unlabeled data and subsequent fine-tuning with a relatively small dataset. Several kinds of BERT models for property prediction have been developed. For instance, Wu *et al*. developed Knowledge-based BERT (K-BERT) pre-trained with three different tasks, predicting properties of individual atom, forecasting overall fingerprints of molecule, and comparing various forms of SMILES representations, thus establishing a comprehensive molecular property prediction framework^[15]^. The average AUC of K-BERT is 0.806 (test set), evaluated on 15 drug discovery-related datasets. The outcome from the ablation experiments of three pre-training tasks has demonstrated that the pre-training could boost the performance of downstream tasks. Xia *et al*. developed MoleBERT that refines the representation of each atom by avoiding simple representations of atomic types^[16]^. It encodes each atom by its surrounding environment and construct a novel vocabulary, which significantly expands the lexicon and mitigates issues stemming from extreme imbalance of atomic types. Moreover, it employs two distinct masking rates during masking, forming a triplet along with the original molecule and training the model through contrastive learning. MoleBERT achieves good performance with an average AUC of 0.740 (test set), evaluated on 8 binary classification tasks. In order to highlight the impact of frequently occurring substructures on molecular property, Li *et al*. developed Functional-Group BERT^[17]^ (FG-BERT), thus converting the task from predicting individual atoms to predicting substructures. It has been validated that FG-BERT performs better than MoleBERT on 8 binary classification tasks (average AUC: 0.7492 for FG-BERT vs. 0.7404 for MoleBERT), and 4 regression tasks (average RMSE: 0.927 for FG-BERT vs. 0.999 for MoleBERT). In order to further validate the performance of the FG-BERT model, experiments with 15 ADMET datasets and 14 breast cancer cell-based phenotypical screening datasets were performed, and the outcome was pretty good, with the average AUC values of 0.813 and 0.856, respectively.

To the best of our knowledge, all the above mentioned models only take the information of molecular two-dimensional structure. In fact, molecules generate bioactivity by binding to its targets and thus three-dimensional conformation are considered more important^[18]^. We hypothesized that the incorporation of spatial information such as bond length and bond angle into the BERT model may further improve model performance^[19]^. Based on this idea, we developed a molecular property prediction model by incorporating bond length and bond angle spatial attention in addition to the usual individual atoms. Through constructing three distinct connection relationships, i.e. atom-atom, atom-bond, and bond-bond, we were able to incorporate the overall topological structure of a molecule into the model so as to enhance attention mechanism in the Transformer architecture. During the attention calculation, we introduced two types of positional information, i.e. bond length, and bond angle. It enables the model to focus on three-dimensional conformation of a molecule during training. To validate its effectiveness, we carried out benchmarking study with 8 public datasets and compared the results with the previously reported state-of-the-art (SOTA) model. Our model, GEO-BERT, demonstrates superior performance across most datasets. Furthermore, we applied GEO-BERT model to real-world activity prediction and drug screening, with DYRK1A-targeted drug discovery as a case study. Top 500 compounds predicted to be active for DYRK1A by the GEO-BERT model were further filtered by conventional computer aided drug design strategy including molecular docking, 20 potential hits were tested for their DYRK1A inhibitory activity. The identification of 2 novel DYRK1A inhibitors (hit rate: 10%) validated the practical applicability of the GEO-BERT model.

## 2. Materials and Methods

### 2.1. Data Collection and Curation

Firstly, approximately 2.13 million molecules were obtained from the ChEMBL^[20]^ (version 30). Like FG-BERT, molecules were selected according to three rules regarding drug-like properties, i.e., molecular weight ≤ 500, ClogP ≤ 5 and number of hydrogen bond donors ≤ 5. A low-energy conformation was generated for each molecule using RDKit (version 2023.03). Since conformers were not successfully generated for approximately 2,000 molecules (mol.GetNumConformers() > 0), these molecules were removed from the data set. Then, bond length and bond angle of each molecule were calculated based on the atom coordinates of each conformation, with the functions implemented in RDKit (i.e., rdkit.Chem.AllChem.EmbedMolecule(mol) and mol.GetConformer().GetPositions(). As a result, a total of 1,454,105 molecules were obtained. They were divided into a training set and a test set at a ratio of 9:1 and ready to use for pre-training.

Eight benchmarks were accessed online and used for transfer learning and comparative analysis, i.e., Tox21, ToxCast, Sider, ClinTox, MUV, HIV, BBBP, and Bace^[21]^.

For the prospective study (i.e. real-world target-based drug discovery practice), the DYRK1A-Dataset that we previously compiled (https://github.com/jwxia2014/DYRK1A-Dataset) was used. It contained 927 molecules collected from ChEMBL and labeled with IC_50_ values. This data set was curated according the following protocol. Following salt removal and standardization, any molecule with more than 20 rotatable bonds and molecular weight greater than 600 was eliminated. The IC_50_ values obtained from multiple bioassays were averaged. The molecules with IC_50_ values below 1 μM were labeled as inhibitors (430), while the others were defined as non-inhibitors (497). This dataset had been randomly divided into two sets at a ratio of 8:2, with data size of the train set (“DYRK1A-train” set) as 742 and the test set (“DYRK1A-test” set) as 185.

### 2.2. The building of GEO-BERT Model

#### 2.2.1. The GEO-BERT Framework

In GEO-BERT, the bond was regarded as an independent token, rather than solely as a channel for information transmission between atoms. As Figure 1A shows, in the stage of pre-training, several atoms and bonds of the molecule were masked. In this way of pre-training, the model learns connectivity between atoms, between bonds as well as between atom and bond.

**Figure 1.**
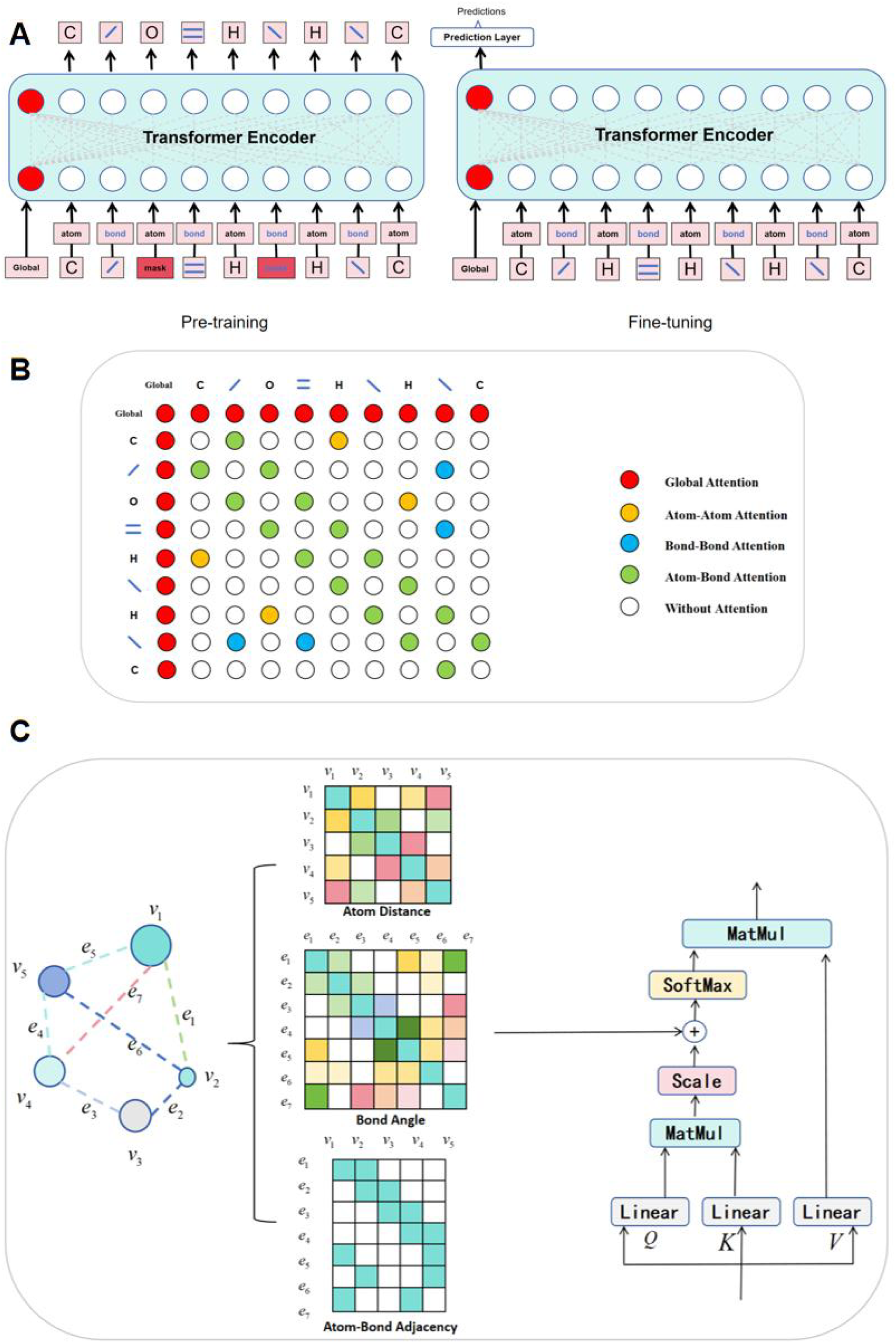
The framework of GEO-BERT. (A)Schematic diagram of pre-training and fine-tuning in the GEO-BERT model. (B) Visualization of adjacency matrices of all atoms and chemical bonds of a molecule during pre-training. According to the connection relationship between atoms and/or chemical bonds of molecules, five types of attention mechanism were defined. (C) Visualization of GEO-BERT’s spatial position matrix, including atomic distance matrix, chemical bond angle matrix, atomic chemical bond adjacency matrix, and the Transformer’s attention calculation process.

In fine-tuning, the pre-trained Global token was extracted and a prediction layer was incorporated to execute molecular property prediction task. As the attention matrix in Figure 1B shows, by leveraging connectivity between atoms, and between atom and bond, as well as between bonds of a molecule, three types of attentions were defined. For all the atoms, attention was exclusively calculated between two atoms linked by a bond. For all the bonds, attention is solely computed between bonds that shared an atom at either end. Likewise, attention is only computed between a bond and each of two atoms on both ends of the bond. With such a definition for connectivity, information was efficiently extracted from the connection between atoms and bonds.

In addition, to calculate spatial position information of a molecule, three additional matrices were incorporated into each attention matrix, as illustrated in Figure 1C. (1) For the atom-atom attention matrix, an Atom Distance matrix was incorporated. It is noted that position information was calculated only for adjacent atoms; for the other atoms, position information was set to 0. (2) For the bond-bond attention matrix, a Bond Angle matrix was incorporated. Similar to Atom Distance matrix, Bond Angle matrix was only calculated between those connected bonds; otherwise, the value in the matrix was set to 0. (3) In the atom-bond attention matrix, as there was no positional information, an Atom-Bond Adjacency matrix was added to balance the above matrices. The attention weight between atoms was affected by distance, i.e., functional relationship between two atoms at different distances varies, aligning with chemical intuition. Similarly, the attention weight between bonds was affected by angle, i.e., functional relationship between two bonds at different angles varies.

To clarify the aforementioned design, the spatial position matrix derived from an exemplified molecule of ethanol is shown in Figure S1. In this figure, it is noted that all the atoms are arranged at the front of the sequence and the bonds at the rear in the figure. In the atom-atom matrix located at the upper left corner, the diagonal entries represent individual atoms. Among the interconnected atoms of the ethanol molecule, the longest distance was observed between the oxygen and one hydrogen atom, while five pairs of C-H atomic distances were equal. Adjacent to C-O (1.4 Å), the shortest distance is between two carbon atoms, i.e. 1.3 Å. For the atom-bond matrix, each bond linked two atoms at both ends, with a maximum of four bonds connecting to carbon atom (C) and only one bond connecting to Hydrogen atom (H). In the bond-bond matrix, the angles formed by three bond pairs, C-O-H, C-C-H, and O-C-H, were slightly greater than the angles formed by the other bonds of the molecule.

In terms of model implementation, the aforementioned three matrices were integrated into the calculation of attention in the Transformer. These matrices were shared by each layer in Transformer, which enabled the model to focus not only on semantic relationship between atoms or bonds, but also on spatial information of the molecule in every attention calculation.

*Color code:

1. Atom Distance: Different colors indicate different distances between atoms, while white indicates no chemical bond between two atoms. Atom distances are symmetrically distributed along the diagonal of the matrix.
2. Bond Angle: Different colors indicate different angles between chemical bonds, while blanks indicate that there is no common atom at the ends of the two bonds. Bond angles are symmetrically distributed along the diagonal of the matrix.
3. Atom-Bond Adjacency: Colors indicate the connection between an atoms and a bond, while blanks indicate that atoms and bonds are not connected.

For the ablation experiment in Section 4.3, to explore the role of different components considered in GEO-BERT, we designed three other different models named as GEO-BERT [a], GEO-BERT [b] and GEO-BERT [c]. GEO-BERT [a] omits the bond length matrix and bond angle matrix. GEO-BERT [b] incorporates the atom distance matrix and bond angle matrix by first multiplying each of them by a learnable parameter and then adding them to the original attention matrix. These learnable parameters are shared across all the layers of GEO-BERT. GEO-BERT [c] employs two special tokens (GLOBAL-ATOM and GLOBAL-BOND) to focus on the atoms and bonds of a molecule respectively, and subsequently concatenates the resulting vectors during fine-tuning. The structural diagrams of GEO-BERT [a], GEO-BERT [b] and GEO-BERT [c] are listed in Figure S2-S4.

#### 2.2.2. The principle of GEO-BERT

The input of the model was a list composed of all the atoms and bonds of a molecule. All the atoms were listed at the front and the bonds were listed behind them, which can be represented as *input* = (*a*_1_, *a*_2_, …, *a_M_*, *b*_1_, *b*_2_, …, *b_N_*). *M* represented the number of atoms, *N* represented the number of bonds.

Based on an embedding matrix *D* ∈ *R^n^*^×*dim*^, the atom and bond vectors in the embedding space were obtained. In the embedding matrix, *n* is the size of the vocabulary composed of all the atoms and bonds existing in all chemical compounds, *dim* is the embedding dimension of each vocabulary, and the input vector is represented as *X* = (*v*_1_, *v*_2_, …, *v_M_*, *e*_1_, *e*_2_, …, *e_N_*).

This process is recorded as formula 1

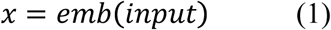

The attention layer in the model contains three matrices *Q*, *K*, and *V*, which are calculated by the formulas 2-5,

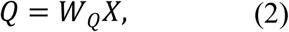

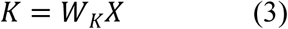

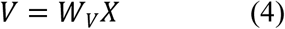

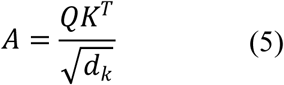

*W_Q_*, *W_K_*, *W_V_* ∈ *R*^d_k_×d_m_^, *d_k_* = *dim*/*H*, *H* is the number of heads with multi-head attention. After Q and K are obtained, the attention A can be calculated.

For any adjacent atom *i* and atom *j* of a molecule, the actual distance between them can be represented using the following formula 6,

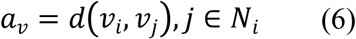

For any adjacent bond *p* and bond *q* of a molecule, the bond angle can be represented using the following formula 7,

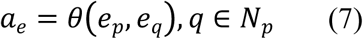

For atom *i* and bond *p* or atom *j* and bond *q* in a molecule, the following formula 8 can be obtained based on whether they are connected in the molecule.

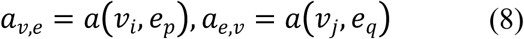

After the above four matrices *a_v_*, *a_e_*, *a_v,e_*, *a_e,v_* were obtained, it was easy to know that *a_v,e_* = *a_v_*_,*e*_^T^, which can form a spatial matrix *A_space_*. The size of the matrix is [*M* + *N*, *M* + *N*], and then it is added to the original attention matrix *A* to obtain a new matrix *A*\*. Finally, the output is calculated by *softmax* and multiplied by *V* to obtain the output value.

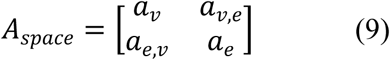

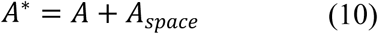

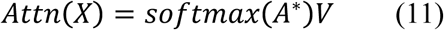

In order to avoid gradient vanishing during calculation process, a residual connection was added to the model after each attention calculation in each layer, and then an *LN* layer was added for layer normalization. The formula is as follows,

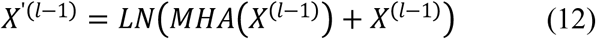

Where *X*^(*l*−1^) represents the intermediate vector of layer *l* − 1, and *X*’^(*l*−1)^ represents the intermediate vector obtained through *LN* in layer *l* − 1.

In order to increase the complexity of each word representation, the feed forward layer (FFN) was added as well. Residual connections were also added during the calculation, and then the layernorm (*LN*) layer was normalized. The formula is as follows:

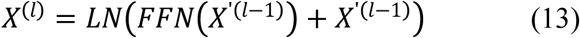

Among them, *X*^(*l*)^ represents the output vector of the *l* − 1 layer obtained through *FFN* and *LN* layers, and serves as the input vector of the *l* layer. The above process fully represents the calculation process of one layer in the Transformer. In our model, a total of six modules (*l*=6) was used.

### 2.3. Pre-training and fine-tuning of GEO-BERT

Following FG-BERT^[17]^, 13 common atoms, i.e., H, C, N, O, S, F, Cl, Br, P, I, Na, B, Se, Si, were defined in molecular vocabulary of GEO-BERT. Rare atoms, e.g. Pb and Hg, were represented as [UNK]. To denote all bonds of a molecule, four symbols (i.e., single, double, triple, benzene) were defined in GEO-BERT. It should be noted that double bonds of benzene and carbon chains were represented individually due to their different physicochemical properties. Lastly, for masked atoms or bonds, [MASK] was introduced. [GLOBAL] was used to gather features from all atoms and bonds in model training. Taken together, the molecular vocabulary defined in GEO-BERT included: [H], [C], [N], [O], [S], [F], [Cl], [Br], [P], [I], [Na], [B], [Se], [Si], [UNK], [single], [double], [triple], [benzene], [MASK], [GLOBAL].

A masking strategy was applied to atoms and bonds during pre-training. All the atoms and bonds were extracted from a molecule and 15% of them were selected. Due to the specific patterns in chemistry, substitution operation was not introduced. Among the selected atoms and bonds, 90% were masked and 10% were unchanged.

1,454,105 molecules from ChEMBL (version 30) were divided into a training set and a test set at a ratio of 9:1. SMILES string representations were converted to molecular graph by RDKit (version 2023.03). For each undirected graph, a global node labeled as Global was introduced, which was connected to all the atoms and bonds of a molecule. During model training, cross-entropy loss was computed respectively for masked atoms and bonds and then aggregated for backpropagation. With a batch gradient-descent algorithm, Adam was used as the optimizer, and the learning rate was set to 1e^-4^. Additionally, GLOBAL-ATOM and GLOBAL-BOND super nodes were included respectively for atoms and bonds. For model validation, the same masking strategy was applied to the test set, and then the overall ratio of correctly predicted atoms and bonds, i.e. Recovery accuracy, was calculated as the metric.

For the public benchmarks, molecules from Tox21, ToxCast, Sider, ClinTox, MUV, HIV, BBBP, and Bace were respectively divided into a training, a validation, and a test set at a ratio of 8:1:1. The public datasets were partitioned using a uniform segmentation approach same as FG-BERT^[17]^, centered on the molecular skeleton division. This method involves grouping molecules into distinct clusters based on their skeletal structure, consequently leading to pronounced distribution disparities between the training and test sets and imposing heightened performance demands on the model.

During the fine-tuning stage, the network layer utilized by the model head for recuperating the masked segment in pre-training step was removed, while a new prediction head composed of two fully-connected layers was incorporated for downstream tasks. In fine-tuning, as the dropout rate and learning rate (lr) significantly affect performance and thus they were optimized during model training. The value of of learning rate was [1e-5, 3e-5 or 5e-5]. The batch sizes were selected from [8, 16, 32, 64, 128]. The dropout rate ranged from 0.1 to 0.5, with an interval of 0.1.

### 2.4. Model Evaluation

For model evaluation, seven metrics were used and defined as formulas (14-20).

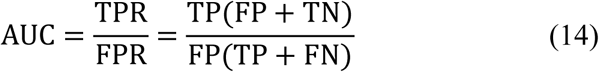

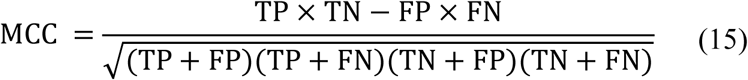

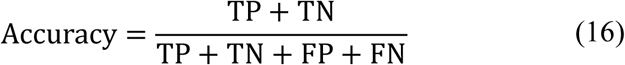

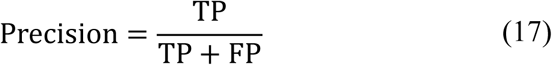

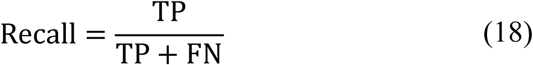

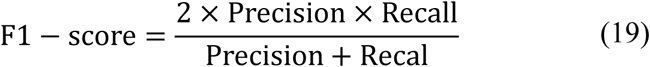

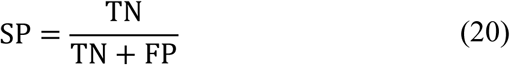

TP represents the number of correctly identified active compounds. TN indicates the number of correctly identified inactive compounds. FP denotes the number of incorrectly identified inactive compounds. FN signifies the number of incorrectly identified active compounds. Based on the aforementioned metrics, more comprehensive metrics such as Accuracy, Precision, Recall, AUC, MCC, F1-Score could be calculated for model evaluation.

### 2.5. Practical Application: DYRK1A inhibitors discovery

As a proof-of-concept, our GEO-BERT model was fine-tuned for dual-specificity tyrosine phosphorylation-regulated kinase 1A (DYRK1A) and applied to discovering DYRK1A inhibitors. DYRK1A plays a crucial role in many physiological processes and can phosphorylate various protein substrates^[22]^. The overexpression of DYRK1A may lead to many diseases such as neurodegenerative diseases, diabetes and cancers. Although many DYRK1A inhibitors have been identified, no drug targeting DYRK1A is approved for clinical use. The discovery of novel DYRK1A inhibitors are still necessary from the perspective of DYRK1A-targeted diseases treatment.

#### 2.5.1. Fine-tuning GEO-BERT model for DYRK1A inhibitory activity prediction

For the DYRK1A dataset, the previously divided “DYRK1A-test” set was still used as the test set (20% of 927 molecules), while the “DYRK1A-train” set was further randomly divided into a smaller train set (60% of 927 molecules) and a validation set (20% of 927 molecules). Fine-tuning was carried out with the train set and evaluated with the validation set and the test set. The number of train epochs was set to 200, with a maximum endurance epoch capped at 30. To be specific, during the training process, if the accuracy based on the validation set did not improve within 30 epochs, the training was stopped. The weights of the model were saved for subsequent evaluation and compound library screening. The test set was used to evaluate the performance of the model with different random seeds, i.e., [0, 1, 2, 3, 4, 5, 6, 7, 8, 9], using the average and variance of all experiments as evaluation metrics. The evaluation metrics included AUC, MCC, Accuracy, Precision, Recall, F1-score, SP.

#### 2.5.2. t-SNE analysis for interpretability of GEO-BERT(DYRK1A)

All the 927 molecules from the DYRK1A dataset were used to analyze the interpretability of the GEO-BERT(DYRK1A) model. The molecules comprised approximately 23,000 atoms and 41,000 bonds and were fed into the GEO-BERT(DYRK1A) model. Consequently, 23,000 atomic vectors and 41,000 bond vectors were generated from the transformer layer. These vectors were then visualized with the t-distributed Stochastic Neighbor Embedding (t-SNE)^[23]^ method.

#### 2.5.3. Uncertainty Assessment of GEO-BERT(DYRK1A)

Three methods, including entropy, dropout, and their ensemble were used to measure uncertainty of the GEO-BERT(DYRK1A) model^[24]^. As for the entropy method, the uncertainty value was calculated by multiplying the model’s prediction probability for each class with the corresponding logarithmic value. As for the dropout method, the dropout operation was adopted, which led to different outcomes during model prediction. Based on the outcomes, the variance was calculated to measure uncertainty. The ensemble method means the averaging of entropy and dropout. GEO-BERT(DYRK1A) model was trained based on the “DYRK1A_train” dataset (including the aforementioned smaller train set and a validation set in Section 2.5.1), while uncertainty was calculated based on the “DYRK1A_test” dataset.

#### 2.5.4. The use of GEO-BERT(DYRK1A) to facilitate virtual screening

After fine-tuning and model evaluation, the GEO-BERT(DYRK1A) model was used for screening the Specs chemical library, and the top 500 compounds were selected based on the predicted activity probability of the model. The compounds were then docked against DYRK1A (PDB code: 7AJ2)^[22]^ that were determined by our group as the structure with the highest ligand enrichment for FRED docking-based virtual screening. Specifically, a maximum of 200 conformations for each compound were generated using OMEGA, and placed into the binding site, evaluated for binding affinity by the Chemgauss4 scoring function, and visually examined for binding modes. Only the compounds that established hydrogen bonds with LEU241 or LYS188 were retained. PAINS-Remover (https://www.cbligand.org/PAINS/) was used to filter the PAINS^[25]^. Lastly, structural clustering based on FCFP_6 fingerprints was carried out with the “Cluster Ligands” module of Discovery Studio (v16.1.0, Dassault Systèmes Biovia Corp.). One or two compounds from each cluster were selected, with priority given to those with greater Chemgauss4 scores as well as experts-based synthetic feasibility.

#### 2.5.5. DYRK1A inhibition assay

The assay was performed in 384-well plates using the ADP-Glo Kinase Assay Kit (Promega). To be specific, this assay measured kinase activity by quantifying the amount of ADP produced during the kinase reaction. The kinase reaction between the kinase and its substrate was performed in an assay buffer consisting of Hepes (Thermo), MgCl_2_ (Sigma), Brij35 (Millipore), EGTA (Sigma), and DTT (MCE). Briefly, ATP and kinase substrate were added to the solution of compounds, kinases and metal ions to initiate the reaction. The reaction was terminated by the addition of ADP-Glo kinase reagent followed by the addition of kinase detection reagent. The mixture was incubated at 25°C. The luminescence was recorded on a microplate reader. Based on the luminescence, the activity values (%) of DYRK1A protein at different concentrations of compounds were determined. GraphPad Prism (GraphPad Software Inc.) was used to determine IC_50_.

## 3. Results

### 3.1. Benchmarking outcome of GEO-BERT on Public Datasets

As illustrated in Table 1, the GEO-BERT model demonstrated the best classification performance, with the highest average AUC (0.7563). This indicates a 0.71% improvement over the suboptimal FG-BERT model. To be specific, the GEO-BERT model performed better than all the previously reported models across 5 out of 8 public datasets, i.e., Tox21 (AUC: 0.791), ToxCast (AUC: 0.672), MUV (AUC: 0.788), HIV (AUC: 0.785), and Bace (AUC: 0.849). Furthermore, its performance also ranked at the 2^nd^ place and was close to FG-BERT model, when it was tested on the Sider dataset and ClinTox dataset. GEO-BERT was also the top-scoring model based on BBBP dataset, with the AUC value greater than 0.70.

**Table 1.**
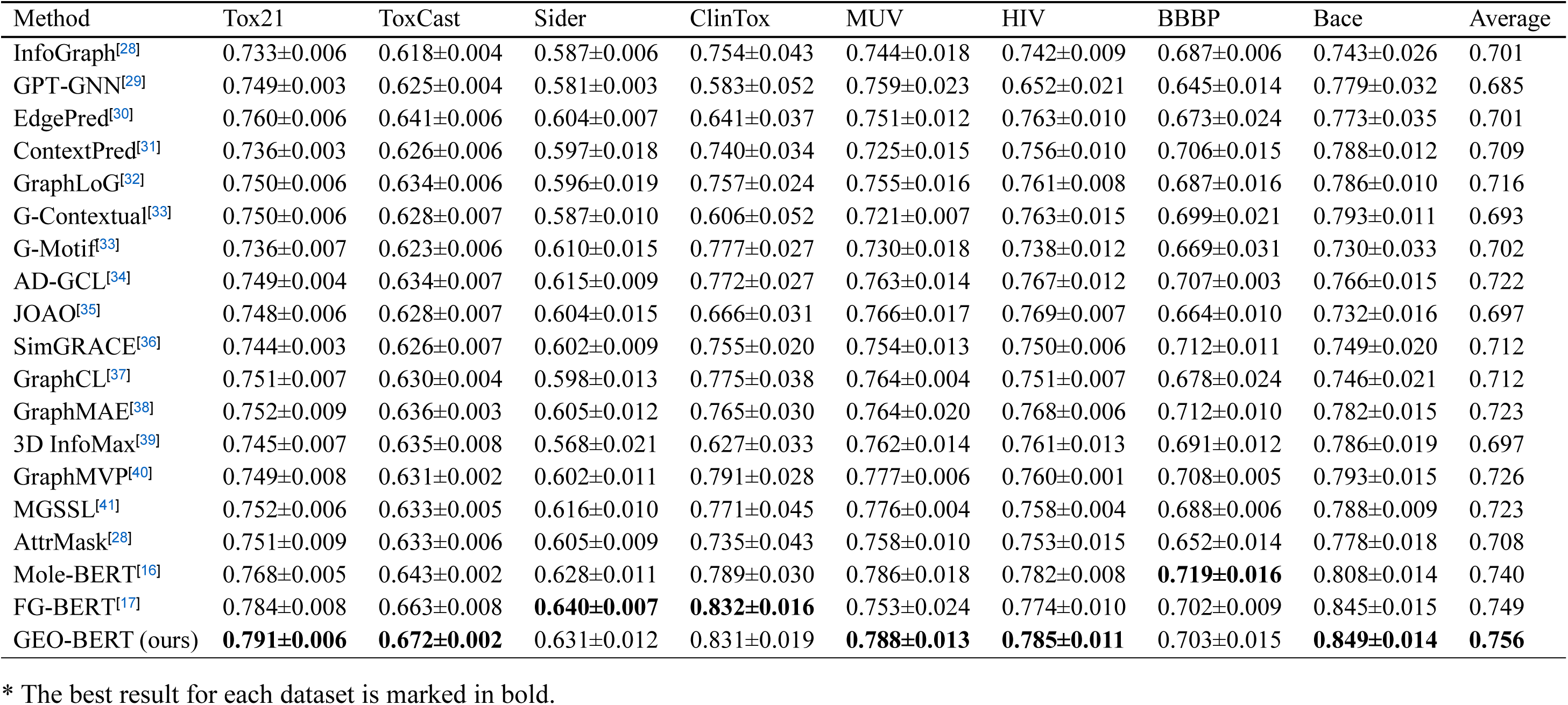
Classification Performance (AUC) of GEO-BERT evaluated based on eight commonly used public datasets.

### 3.2. Performance of GEO-BERT for DYRK1A inhibitory activity prediction

We built GEO-BERT model for DYRK1A by transfer learning with the pre-trained GEO-BERT model and the DYRK1A-dataset. Firstly, the weights of the pre-trained GEO-BERT model based on 1,454,105 molecules from ChEMBL were saved, and then the DYRK1A-dataset was used to fine-tune the model. The classification performance of GEO-BERT(DYRK1A) is shown in Table 2. It achieved good performance with AUC being 0.875 and MCC being 0.610.

**Table 2.** Classification Performance of GEO-BERT (DYRK1A), evaluated on the DYRK1A dataset.

| Method | AUC | MCC | Accuracy | Precision | Recall | F1-score | SP |
| --- | --- | --- | --- | --- | --- | --- | --- |
| RF | 0.821±0.007 | 0.535±0.013 | 0.768±0.018 | 0.770±0.007 | 0.744±0.036 | 0.755±0.009 | 0.790±0.012 |
| SVM | 0.746±0.011 | 0.339±0.042 | 0.665±0.006 | 0.630±0.018 | 0.756±0.012 | 0.677±0.017 | 0.579±0.009 |
| XGBoost | 0.816±0.009 | 0.448±0.007 | 0.724±0.007 | 0.724±0.009 | 0.700±0.017 | 0.714±0.007 | 0.747±0.007 |
| KNN | 0.774±0.014 | 0.372±0.014 | 0.687±0.025 | 0.682±0.021 | 0.667±0.031 | 0.673±0.020 | 0.705±0.025 |
| DNN | 0.831±0.006 | 0.492±0.006 | 0.746±0.014 | 0.759±0.007 | 0.700±0.006 | 0.731±0.006 | 0.790±0.008 |
| Chemprop | 0.794±0.008 | 0.460±0.018 | 0.730±0.019 | 0.717±0.011 | 0.733±0.014 | 0.722±0.012 | 0.726±0.011 |
| MG-BERT | 0.841±0.005 | 0.515±0.013 | 0.757±0.009 | 0.785±0.018 | 0.689±0.008 | 0.732±0.009 | 0.821±0.015 |
| FG-BERT | 0.852±0.009 | 0.556±0.008 | 0.778±0.011 | 0.775±0.009 | 0.767±0.013 | 0.771±0.011 | 0.790±0.007 |
| GEO-BERT (ours) | **0.875±0.014** | **0.610±0.014** | **0.805±0.008** | **0.807±0.012** | **0.811±0.009** | **0.808±0.013** | **0.821±0.007** |

\* The best result for DYRK1A dataset is marked in bold.

To explore whether GEO-BERT(DYRK1A) had advantages, we built models with five classical machine learning algorithms, i.e. RF, SVM, XGBoost, KNN, and DNN, and the classical fingerprints, i.e., MACCS to represent molecules. In addition, we trained models with a well-known supervised deep learning approach, Chemprop^[26]^ and other two BERT models, MG-BERT^[27]^ and FG-BERT^[17]^.

The data sets used for training and evaluating the nine models mentioned above remained consistent with those for GEO-BERT(DYRK1A) (cf. Section 2.5.1). Among all the nine models, GEO-BERT(DYRK1A) showed the best performance. Figure 2 reveals that, only FG-BERT and our GEO-BERT had AUC over 0.85, which were much greater than the classical machine learning models as well as the graph-based model, Chemprop. In addition, only our GEO-BERT model had MCC exceeding 0.6. The data indicated that GEO-BERT is the best model for DYRK1A inhibitory activity prediction.

**Figure 2.**
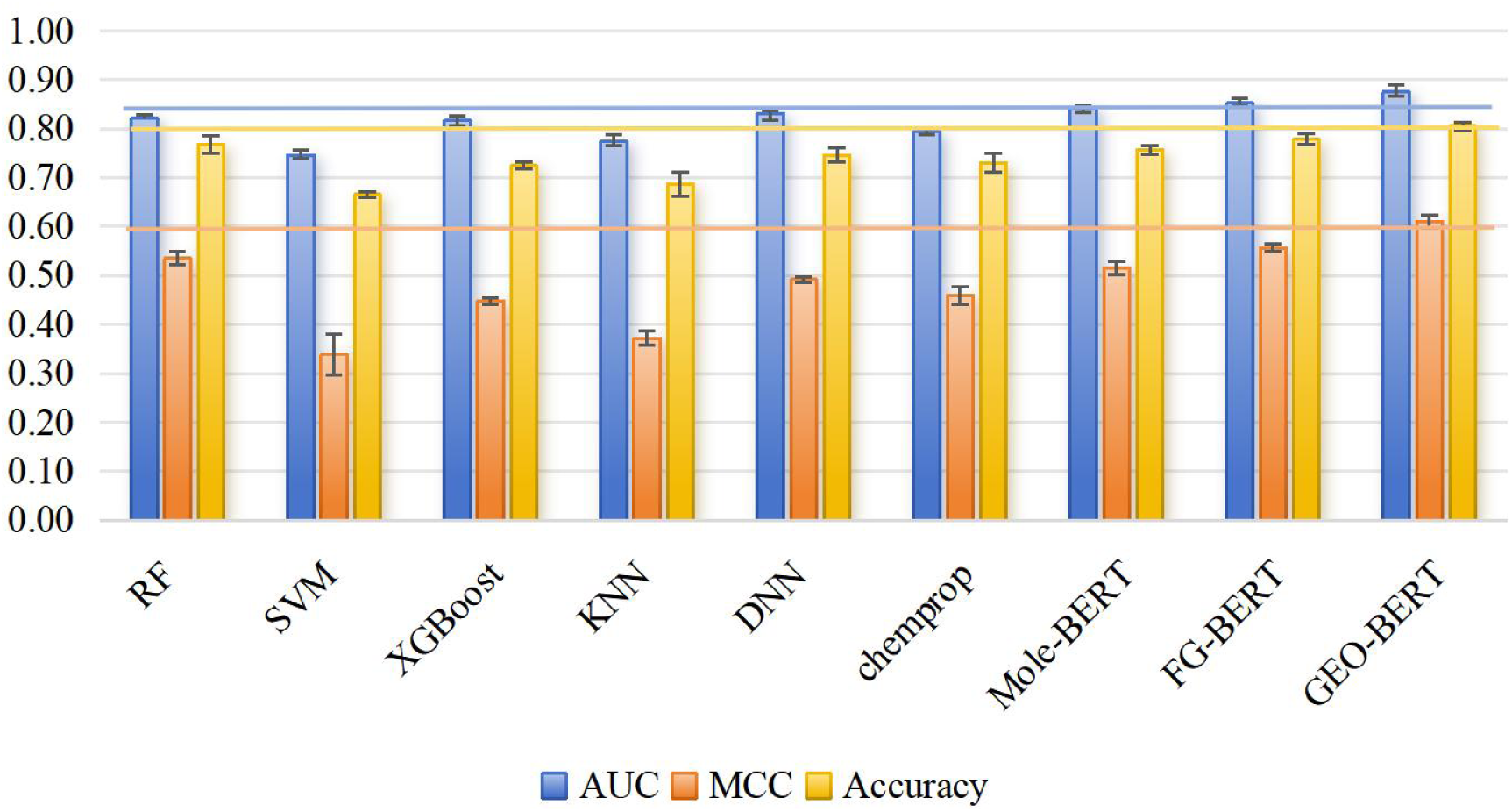
Classification performance of GEO-BERT(DYRK1A) and compassion with other models for prediction of DYRK1A inhibitory activity. Blue line, AUC as 0.85; orange line, MCC as 0.6; yellow line, Accuracy as 0.8.

### 3.3. Interpretability of GEO-BERT(DYRK1A) based on t-SNE analysis

927 molecules from the DYRK1A dataset comprised approximately 23,000 atoms and 41,000 bonds. These data were fed into the GEO-BERT(DYRK1A) model. As a result, 23,000 atomic vectors and 41,000 bond vectors were generated from the transformer layer. These vectors were then visualized with the t-distributed Stochastic Neighbor Embedding (t-SNE)^[23]^ method. Notably, the majority of atoms are carbon atoms, and the most prevalent types of chemical bonds are single and aromatic bonds, with almost no triple bonds. As Figure 3A and Figure 3B show, in terms of atoms and chemical bonds, GEO-BERT(DYRK1A) performed well at distinguishing various types of feature information.

**Figure 3.**
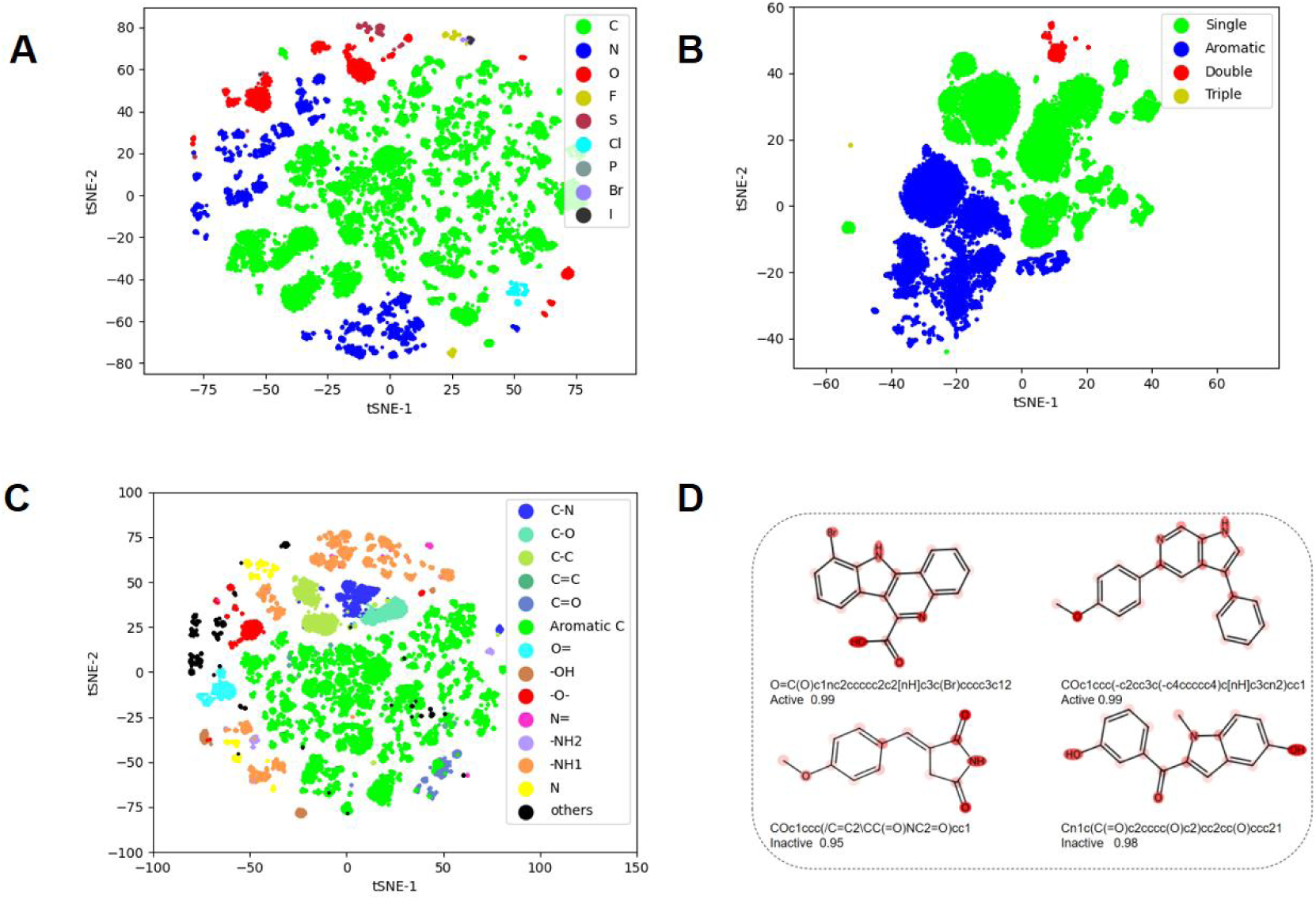
Interpretability analysis of GEO-BERT(DYRK1A). (A) Visualization of t-SNE for embedding vectors of different atom types. B) Visualization of t-SNE for embedding vectors of different bond types. (C)Visualization of t-SNE for embedding vectors of different atomic clusters. (D) Visualization of the attention of representative molecules of the DYRK1A dataset, with darker colors indicating greater attention weights.

In the t-SNE representation of atoms, we observed significant distribution discrepancies even for the atoms of the same type. Consequently, leveraging the surrounding atom information of each atom, we further segmented them into distinct functional groups, reevaluating model performance through t-SNE analysis. Figure 3C illustrates the model’s robustness at identifying different atomic clusters, with consistent aggregation observed among the identical functional groups from different molecules. The identification of highly frequent functional groups by the GEO-BERT(DYRK1A) model might be useful for future molecular design.

Furthermore, we selected two representative active and two inactive molecules from the DYRK1A dataset and visualized their attention weights in the attention head of GEO-BERT(DYRK1A), as depicted in Figure 3D. In Natural Language Processing, the attention weights of an attention head in the Transformer model can reflect the importance of each word in the vocabulary list. This definition could also be applicable to molecules. In two active molecules, the high-attention region represents the substructures that contribute to the activity, while in those inactive molecules, the high-attention region may represent the substructures that impaired the activity. These relationships between the substructure and activity of compounds uncovered by the model may be applied to structural optimization of DYRK1A inhibitors.

### 3.4. Uncertainty of GEO-BERT(DYRK1A)

As shown in Figure 4A, we sequentially removed the samples of high uncertainty ranking at the top 10% of the “DYRK1A_test” dataset and calculated MCC for the remaining data. As the uncertainty index decreased, the model performance improved. Although there was a significant decrease in MCC when the uncertainty decreased from 0.3 to 0.2, the value of MCC still remained around 0.5. The MCC based on other uncertainty metrics is shown in Figure S5. As shown in Figure 4B, (1) the distribution of four types of samples (TP, TN, FP, FN) varied along with different uncertainty values. To be specific, as the uncertainty value increased, the number and proportion of FP and FN also increased. It indicates the prediction accuracy of GEO-BERT(DYRK1A) decreased with the increase of uncertainty. Nevertheless, GEO-BERT(DYRK1A) predicted accurately when the uncertainty value was low. (2) Most samples in the “DYRK1A_test” data set were distributed in the areas with moderate uncertainty values (0.4-0.6), and there were a considerable proportion of samples when the uncertainty value is 0. The result based on other uncertainty metrics can be seen in Figure S6.

**Figure 4.**
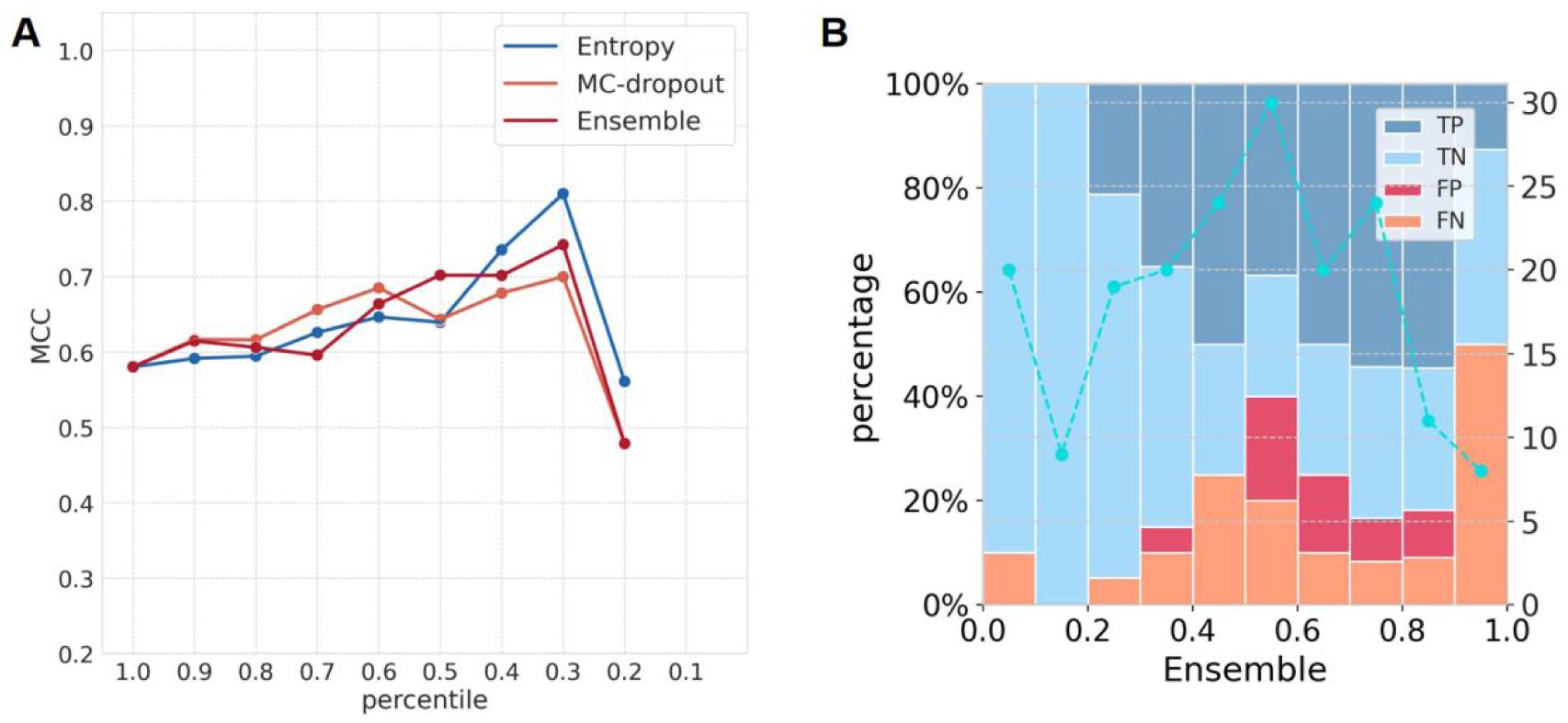
Uncertainty analysis of GEO-BERT(DYRK1A), based on three uncertainty metrics and the “DYRK1A_test” dataset. (A) The variation of MCC in GEO-BERT(DYRK1A) when the uncertainty value changes, where the uncertainty values are calculated by entropy, dropout, and ensemble methods, respectively. (B) The distribution of various types of samples (TP, TN, FP, FN) predicted by GEO-BERT(DYRK1A) within different uncertainty intervals based on Ensemble method. The horizontal axis represents the uncertainty interval, and the vertical axis represents the total number of samples (blue dashed line) and the respective percentage of samples (TP, TN, FP, FN; histogram) within each uncertainty interval.

### 3.5. Hit compounds identified by the computational workflow incorporating GEO-BERT (DYRK1A)

Firstly, we used the GEO-BERT (DYRK1A) model to predict DYRK1A inhibitory activity of Specs compounds (>200K), and then ranked them according to their probability of being active. Top 500 compounds were selected for further screening with conventional structure-based approaches, i.e. molecular docking. From molecular docking against the protein structure (PDB code: 7AJ2), which had been validated as an optimal structure for DYRK1A inhibitors discovery^[22]^, we selected 344 compounds with FRED Chemgauss4 scores less than −10. According to the criteria based on the binding modes, we further narrowed down the number of potential hits to 119. However, only 114 compounds passed the PAINS-Remover. By structural clustering of these compounds based on FCFP_6 fingerprints, we picked 20 diverse compounds. Their chemical structures, FRED Chemgauss4 scores and probability values of being active are listed in Table S3. We tested them for their DYRK1A inhibitory activity at the concentration of 10 μM. As shown in Table S9, compound **X1**, **X6**, **X11** and **X17** had the inhibition rate greater than 50% for DYRK1A. Their IC_50_ values were further determined based on the dose-response curves (cf. Figure S7). Among them, the IC_50_ values of **X11** and **X17** were 0.30 μM and 0.55 μM, respectively, indicating that they were potent DYRK1A inhibitors (<1 μM). The predicted binding modes of **X11, X17** to DYRK1A are shown in Figure 5. Compound **X11** formed hydrogen bonds with two key amino acid residues GLU239/LEU241, and hydrophobic interactions with VAL173/ALA186/LEU294/VAL306. Unlikely, compound **X17** interacted with LYS188 via a hydrogen bond, and formed hydrophobic interactions with ILE165/VAL222/VAL306. To explore chemical novelty of these inhibitors, we calculated structural similarity between each of the two hits and the known DYRK1A inhibitors that we collected from ChEMBL33 (https://www.ebi.ac.uk/chembl/, accessed May. 2023). As shown in Figure S7, based on FCFP_6 fingerprints, the maximum structural similarity (Tanimoto coefficient) was only 0.165 to **X11**, and 0.267 for **X17**. It demonstrated that those two potent hits identified by GEO-BERT(DYRK1A)-based computational workflow were structurally novel.

**Figure 5.**
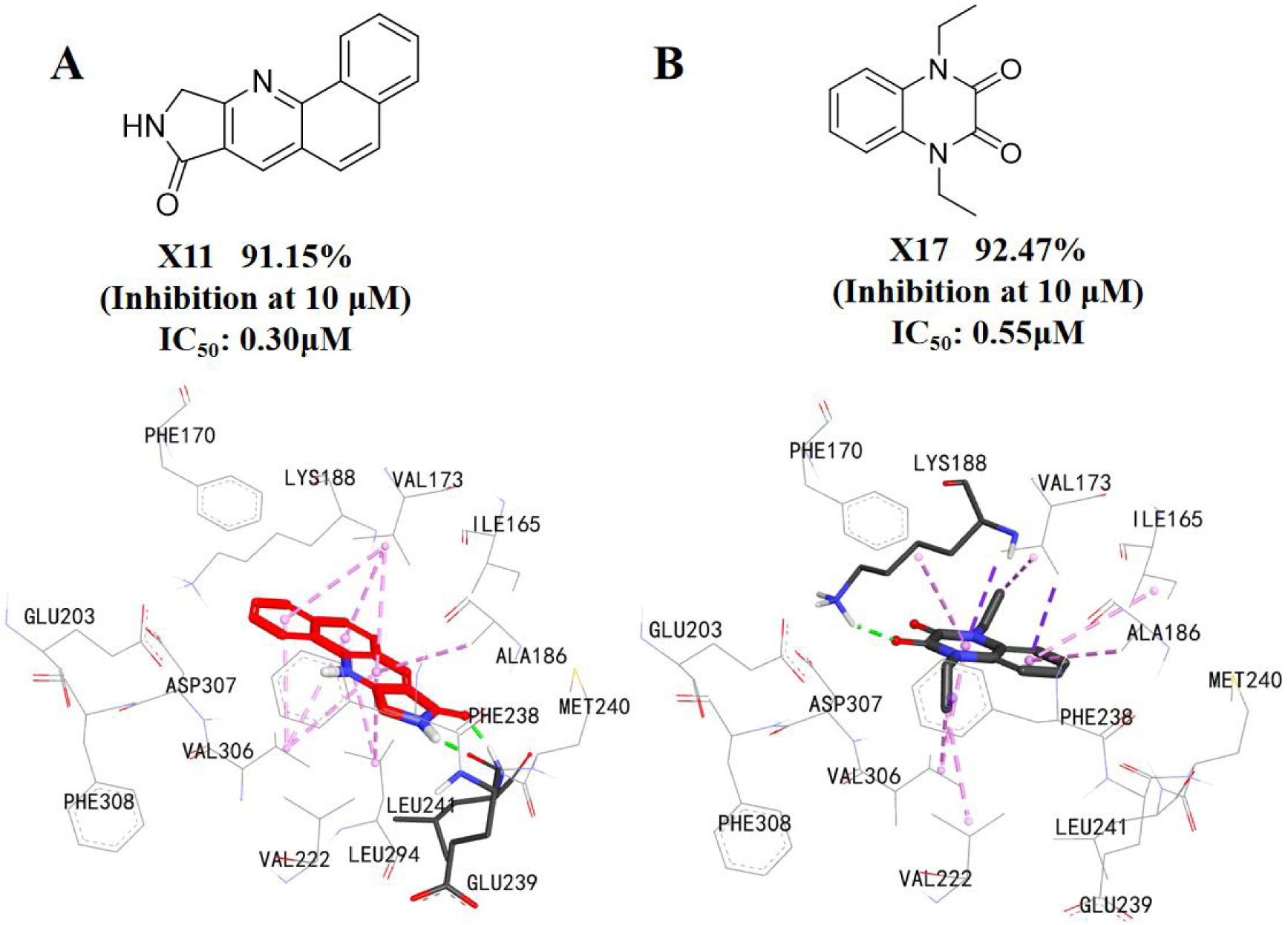
The hit compounds (A) and (B) identified by virtual screening facilitated by GEO-BERT(DYRK1A) and the predicted binding mode to DYRK1A.

In addition, we used the t-SNE method to show the chemical space of the molecules from the “DYRK1A_test” dataset. The feature vectors corresponding to each sample in the last layer of GEO-BERT(DYRK1A) were extracted and visualized. It can be seen from Figure 6 that TN and FN samples were concentrated together, while TP and FP were close in distance, showing good predictive ability of the model. Based on this plot, we further analyzed the chemical space of those two experimentally validated hits **X11** and **X17**. It can be seen that they have lower uncertainty values, and their distributions in the chemical space were relatively close but distant from the molecules of the “DYRK1A_test” dataset.

**Figure 6.**
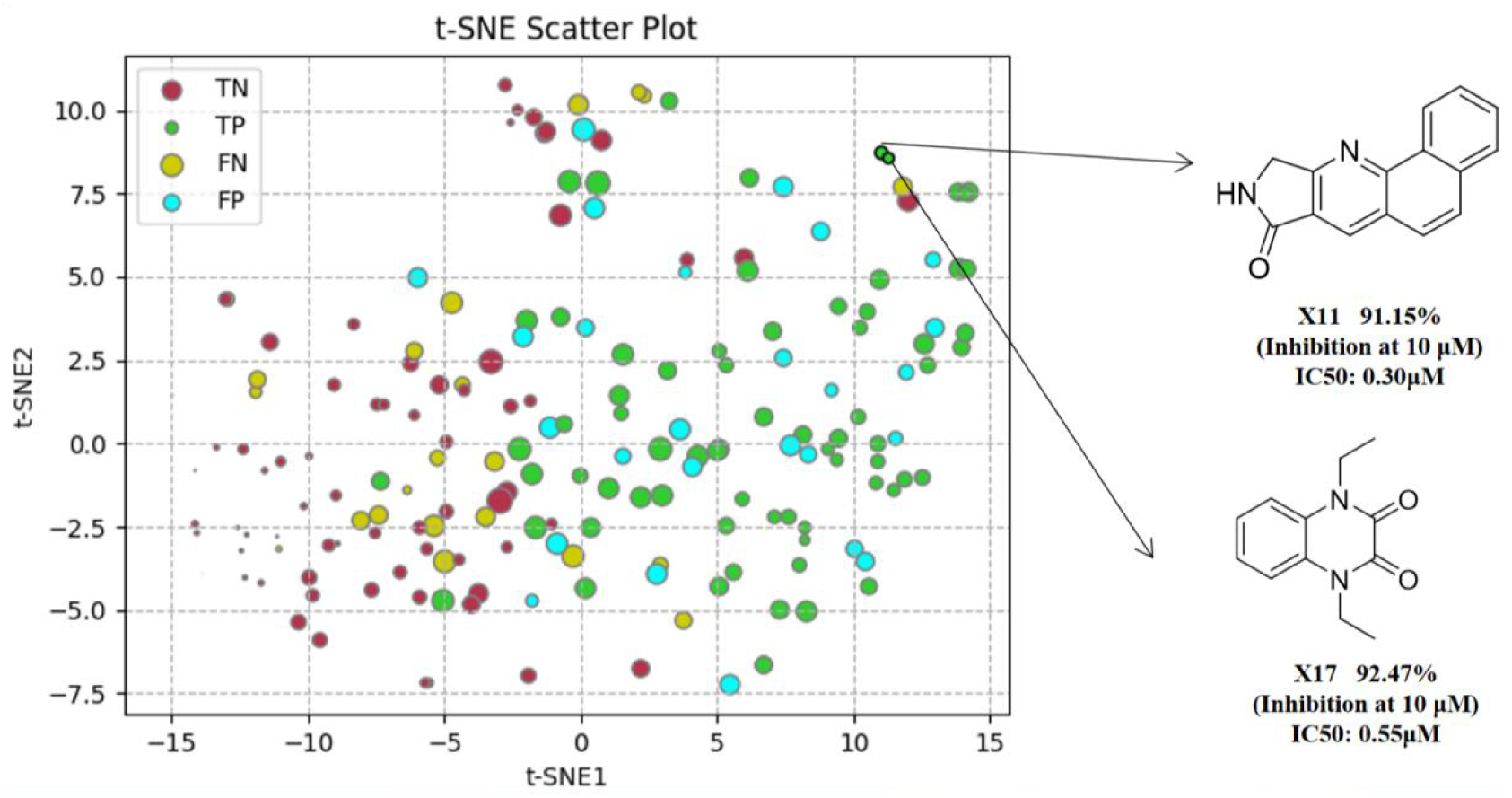
The distribution of four types of samples (TP, TN, FP, and FN) in the GEO-BERT model on the “DYRK1A_test” dataset. Visualization of chemical space using the t-SNE method, where points of different colors represent four types of samples, and the size of the points represents the uncertainty value of each sample, calculated by the ensemble method. Two validated hit compounds **X11** and **X17** are marked separately in the feature space.

## 4. Discussions

### 4.1. The effect of scale parameters on GEO-BERT(DYRK1A)

We explored the effects of different combinations of configurations (Layers, Heads, Embedding size and FFN size) on model performance of GEO-BERT(DYRK1A). In order to compare their performance, we followed the settings of MG-BERT^[27]^ and designed three models with different scale parameters, i.e. GEO-BERT-[SMALL], GEO-BERT-[MEDIUM] and GEO-BERT-[LARGE]. The number of layers was 3, 6, and 12, respectively. In order to match the size of the layers, their Heads, embedding size, and FFN size also adjusted accordingly. The AUC of GEO-BERT(DYRK1A) were utilized as metrics, with the results presented in Table 3.

**Table 3.** Comparison of three GEO-BERT(DYRK1A) models with different configurations.

| Name | Layers | Heads | Embedding size | FFN size | AUC |
| --- | --- | --- | --- | --- | --- |
| GEO-BERT-[SMALL] | 3 | 2 | 128 | 256 | 0.853 |
| GEO-BERT-[MEDIUM] | 6 | 4 | 256 | 512 | 0.875 |
| GEO-BERT-[LARGE] | 12 | 8 | 576 | 1152 | 0.865 |

As shown in the table, GEO-BERT-[MEDIUM] performed the best among the three models, with the AUC value of 0.8750, respectively. When the scale parameters were too small, it may fail to capture sufficient information from the molecules. Conversely, an excessive number of layers may lead to model overfitting and decreased AUC. Accordingly, we chose the medium size to balance model complexity and model performance. The ROC curve for GEO-BERT with parameters of different scales can be seen in Figure S8(A).

### 4.2. Ablation experiment of GEO-BERT(DYRK1A)

To explore the role of spatial information considered in GEO-BERT, we designed GEO-BERT [a], GEO-BERT [b] and GEO-BERT [c]. The details of these models are described in Section 2.2.1. As depicted in Table 4, the absence of atom distance matrix and bond angle matrix in GEO-BERT[a] resulted in decreased performance (AUC: 0.860 vs. AUC: 0.875; MCC: 0.600 vs. MCC: 0.610), which indicated the importance of spatial information. Furthermore, the introduction of two learnable parameters that multiplied atom distance matrix and bond angle matrix in GEO-BERT [b] also impaired model performance, when compared with the direct incorporation of fixed distance and angle values in GEO-BERT (AUC: 0.845 vs. AUC: 0.875 / MCC: 0.535 vs. MCC: 0.610). GEO-BERT [c] partitioned atoms and bonds by splitting a molecule into two segments. However, since a molecule is a unified chemical entity comprising both atoms and bonds, GEO-BERT that employed the single GLOBAL mechanism yielded better performance than GEO-BERT[c] (AUC: 0.864 vs. AUC: 0.875; MCC: 0.549 vs. MCC: 0.610). The ROC curves in Figure S8(B) demonstrate consistent results.

**Table 4.** Classification Performance of various variants of GEO-BERT(DYRK1A) for DYRK1A inhibitory activity prediction.

| Models | AUC | MCC | Accuracy | Precision | Recall | F1-score | SP |
| --- | --- | --- | --- | --- | --- | --- | --- |
| GEO-BERT[a] | 0.860±0.006 | 0.600±0.015 | 0.800±0.009 | 0.802±0.009 | 0.767±0.013 | 0.788±0.007 | 0.812±0.021 |
| GEO-BERT[b] | 0.845±0.015 | 0.535±0.007 | 0.768±0.011 | 0.759±0.007 | 0.767±0.012 | 0.761±0.015 | 0.768±0.014 |
| GEO-BERT[c] | 0.864±0.009 | 0.549±0.008 | 0.773±0.015 | 0.745±0.031 | 0.789±0.006 | 0.769±0.007 | 0.737±0.011 |
| GEO-BERT | **0.875±0.014** | **0.610±0.014** | **0.805±0.008** | **0.807±0.012** | **0.811±0.009** | **0.808±0.013** | **0.821±0.007** |

\* The best result for DYRK1A dataset is marked in bold.

### 4.3. Performance of GEO-BERT(DYRK1A) on out of distribution data

Generalization is a pretty important metric of machine learning models, as it somewhat determines model performance in real-world practical application. In this study, we used the following method to evaluate generalization of our GEO-BERT(DYRK1A) model. Firstly, we calculated pairwise similarity in terms of Tanimoto coefficient between molecules of the “DYRK1A-test” set and those of the “DYRK1A-train” set using MACCS fingerprints. Any molecule of the “DYRK1A-test” set with the similarity value above a specified threshold was excluded, and the model was then evaluated on the revised test set. As the threshold was set to 1.0, 0.95, 0.90, 0.85, 0.80 and 0.75, respectively, the performance in terms of AUC and MCC was recorded for each threshold.

As depicted in Figure 7, as the molecular similarity decreased, i.e. the molecules were more dissimilar to those used for model training, the performance of GEO-BERT(DYRK1A) also gradually declined. Nevertheless, AUC still remained above 0.6, when the similarity threshold was 0.75. In addition, we chose FG-BERT, which is also based on the transformer infrastructure, as a comparison. From the figure, it can be seen that GEO-BERT(DYRK1A) outperformed FG-BERT(DYRK1A) in AUC and MCC at various thresholds. In summary, GEO-BERT(DYRK1A) demonstrated robust performance when encountering diverse molecules.

**Figure 7.**
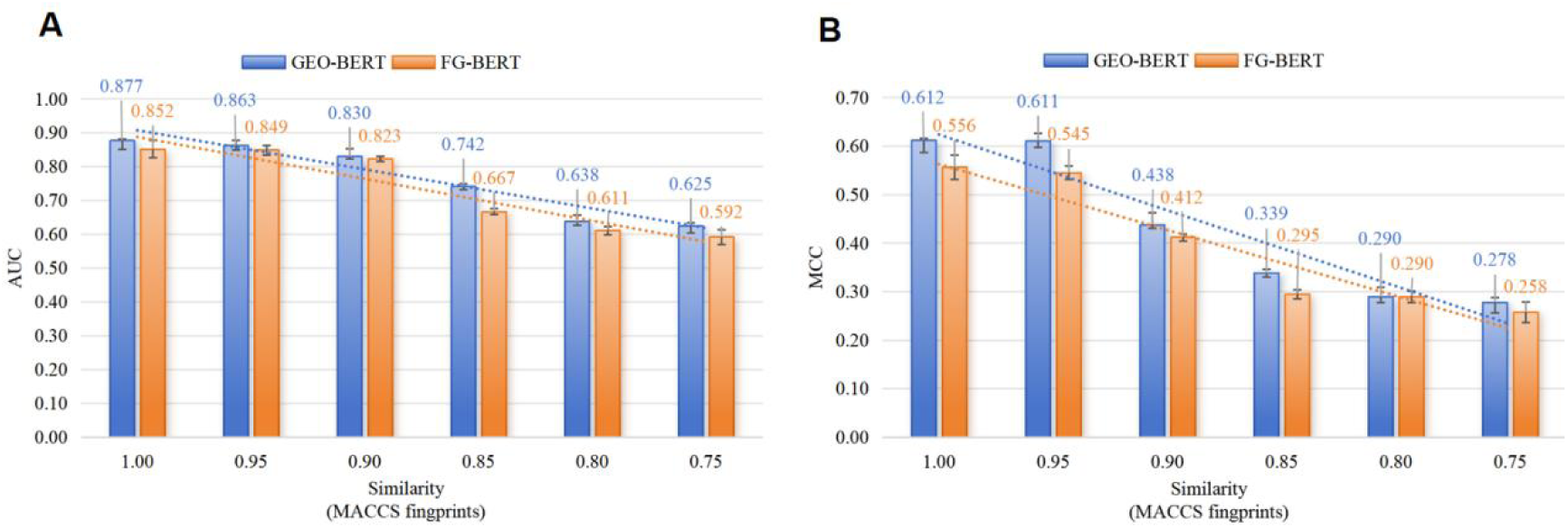
Comparison between the GEO-BERT(DYRK1A) model and the FG-BERT(DYRK1A) model in terms of generalization. The x axis is the similarity threshold used to exclude similar molecules from the “DYRK1A-test” set, while the y axis is the metrics used in the comparison. (A) Comparison in the metric of AUC. (B) Comparison in the metric of MCC.

## 5. Conclusion

In this study, we propose a GEO-BERT model that uses three-dimensional features of molecular conformation as additional information to commonly used molecular two-dimensional structure. Though previous models such as GraphMVP^[40]^ also used three-dimensional information, our GEO-BERT has the following technical advantages. (1) GraphMVP and other models are still based on graph neural networks, which cannot avoid the inherent drawbacks of overfitting and oversmooth caused by the message passing mechanism of graph neural networks. By contrast, GEO-BERT integrates graph message passing into Transformer and solves this problem by using attention mechanism. (2) In graph neural networks, the information of chemical bonds is added to the surrounding atomic features through aggregation operations in message passing. The subsequent graph pooling layer only uses features from atoms, while GEO-BERT can directly utilize information from chemical bonds by treating them as independent tokens. (3) For the first time, this work uses bidirectional autoencoder to study molecules with three-dimensional information. By predicting the masks of atoms and chemical bonds, the model focuses on the three-dimensional spatial environment inside the molecule. (4) More importantly, GEO-BERT was applied to real-world drug discovery, and facilitated the discovery of two novel and potent DYRK1A inhibitors at a high hit rate, demonstrating its high practical utility. It should be noted that all the previous models have never been experimentally validated.

In summary, we have developed a technically novel and practically effective Geometry-based BERT model for molecular property prediction. Due to the open-source feature, it is anticipated that this method will have a wide application in early-stage drug discovery.

## 6. ASSOCIATED CONTENT

### 6.1. Data and Software Availability

The source code is available at the open-source GitHub repository (https://github.com/drug-designer/GEO-BERT), where all the parameters of the trained models and the curated datasets that can be used for prospective study are also provided.

### 6.2. Supplementary materials

Supplementary material associated with this article can be found, in the online version, at www.###.

