## Supporting_Information for "Geometry-based BERT: an experimentally validated deep learning model for molecular property prediction in drug discovery"

**Table of Contents**

**Table S1.** Comparison of feature information used by GEO-BERT and other model. P3

**Table S2.** The details of 8 public benchmarks.	P4

**Table S3.** 20 potential hits identified by GEO-BERT(DYRK1A) and structure-based virtual screening and tested for bioactivity against DYRK1A (%Inhibition at 10 μM). P5

**Figure S1.** Visualization of the three-dimensional spatial position matrix.	P7

**Figure S2.** GEO-BERT [a] omits the Atom Distance and Bond Angle matrix, using a 0-1 mask matrix instead. P8

**Figure S3.** GEO-BERT [b] incorporates the atom distance matrix and bond angle matrix by first multiplying each of them by a learnable parameter and then adding them to the original attention matrix. These learnable parameters are shared across all the layers of GEO-BERT. P9

**Figure S4**. GEO-BERT [c] employs two special tokens (GLOBAL-ATOM and GLOBAL-BOND) to focus on the atoms and bonds in the molecule respectively, and subsequently concatenates the resultant vectors during the fine-tuning phase. P10

**Figure S5.** Uncertainty analysis of GEO-BERT(DYRK1A), based on other six uncertainty metrics and the “DYRK1A_test” dataset.	P11

**Figure S6.** Distribution of various types of samples in the GEO-BERT model under six uncertainty metrics.	P12

**Figure S7.** The inhibitory activity and structural novelty of compound **X1** (a), **X6** (b), **X11** (c) and **X17** (d), with Harmine as the reference compound (IC_50_: 21.24nM). P13

**Figure S8.** ROC curves of GEO-BERT on the DYRK1A dataset. P14

**Table S1.** Comparison of feature information used by GEO-BERT and other models

| **Models** | **Bidirectional autoencoder model** | **Use information from atoms** | **Use information from functional groups** | **Use information from chemical bond** | **Use geometric structure information of molecules** |
| --- | --- | --- | --- | --- | --- |
| MG-BERT | √ | √ | × | × | × |
| FG-BERT | √ | √ | √ | × | × |
| GEO-BERT | √ | √ | × | √ | √ |

**Table S2.** The details of 8 public benchmarks.

| Datasets | Type | Tasks | Molecules | Metrics |
| --- | --- | --- | --- | --- |
| BACE | Classification | 1 | 1513 | ROC-AUC |
| BBBP | Classification | 1 | 2039 | ROC-AUC |
| ClinTox | Classification | 2 | 1478 | ROC-AUC |
| SIDER | Classification | 27 | 1427 | ROC-AUC |
| Tox21 | Classification | 12 | 7831 | ROC-AUC |
| HIV | Classification | 1 | 41127 | ROC-AUC |
| MUV | Classification | 17 | 93087 | ROC-AUC |
| ToxCast | Classification | 617 | 8575 | ROC-AUC |

BACE: binary classifications for a collection of human BACE-1 inhibitor compounds.

BBBP: binary classifications of compounds based on their blood-brain barrier permeability characteristics.

ClinTox: qualitative data on drugs that have received FDA approval and drugs that did not pass clinical trials due to toxicity concerns.

SIDER: categorize drug side effects into 27 classes based on the affected organ systems.

Tox21: measurements of compound toxicity on 12 distinct targets, such as nuclear receptors and stress response pathways, using qualitative methods.

HIV: assign binary indicators to compounds based on their capacity to inhibit HIV replication.

MUV: a refined nearest neighbor analysis was employed to select a subset from PubChem BioAssay.

ToxCast: in vitro high-throughput screening was utilized to generate toxicology data for a vast collection of compounds.

**Table S3.** 20 potential hits identified by GEO-BERT(DYRK1A) and structure-based virtual screening and tested for bioactivity against DYRK1A (%Inhibition at 10 μM).

| ID | Specs  IDNUMBER | Chemical tructure | pred | FRED Chemgauss4 score  (FRED) | %Inhibitionc  mean  (10 μM) |
| --- | --- | --- | --- | --- | --- |
| X1 | AI-942/13331060 | 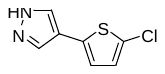 | 0.9955511 | -11.781 | 52.46 |
| X2 | AK-918/43220126 | 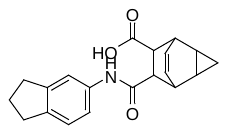 | 0.99075705 | -11.1458 | 24.99 |
| X3 | CS-004/03987021 | 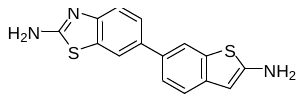 | 0.9927395 | -11.5559 | 47.50 |
| X4 | AF-399/15128324 | 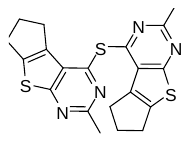 | 0.9903137 | -15.3754 | 47.41 |
| X5 | AN-652/37302001 | 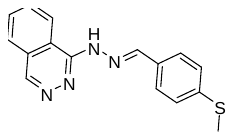 | 0.9892278 | -12.3364 | 34.13 |
| X6 | AC-907/25005013 | 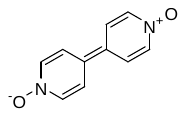 | 0.9941691 | -10.7204 | 52.84 |
| X7 | AG-690/12245813 | 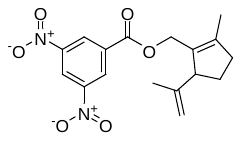 | 0.9900766 | -12.6243 | 10.29 |
| X8 | AO-365/43473525 | 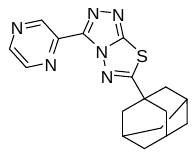 | 0.9897356 | -10.7742 | 8.44 |
| X9 | AK-968/41924802 | 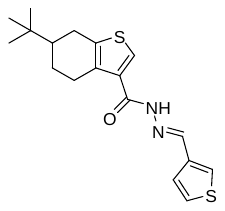 | 0.99033 | -12.2127 | 23.14 |
| X10 | AN-056/25013026 | 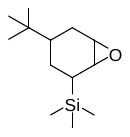 | 0.9945117 | -10.6502 | 38.49 |
| X11 | AP-355/43470180 | 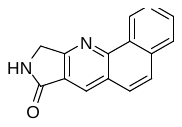 | 0.9915072 | -12.1996 | 91.15 |
| X12 | AK-918/12943318 | 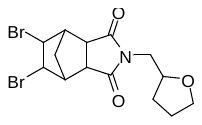 | 0.9913954 | -11.9064 | 41.45 |
| X13 | AT-057/43469674 | 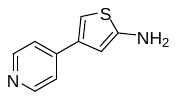 | 0.9899671 | -11.0625 | 25.64 |
| X14 | AG-690/33046024 | 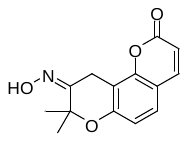 | 0.99374926 | -10.9288 | 38.58 |
| X15 | AE-848/34401046 | 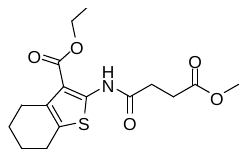 | 0.9905151 | -12.0988 | 18.29 |
| X16 | AF-399/40654548 | 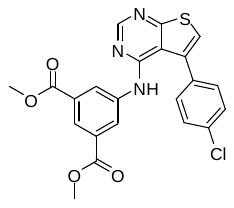 | 0.98941934 | -13.4615 | 29.77 |
| X17 | AO-080/43378446 | 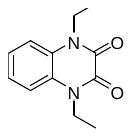 | 0.9902102 | -10.0861 | 92.47 |
| X18 | AN-329/41104307 | 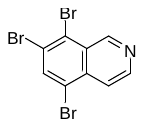 | 0.9904659 | -12.0653 | 41.50 |
| X19 | AK-968/41924692 | 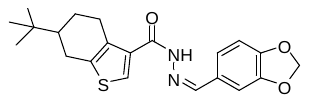 | 0.9904399 | -12.0704 | 12.42 |
| X20 | AG-205/36623038 | 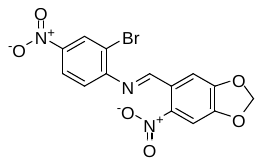 | 0.98967034 | -10.0968 | 29.31 |
| / | Harmine |  | / | / | 98.79 |

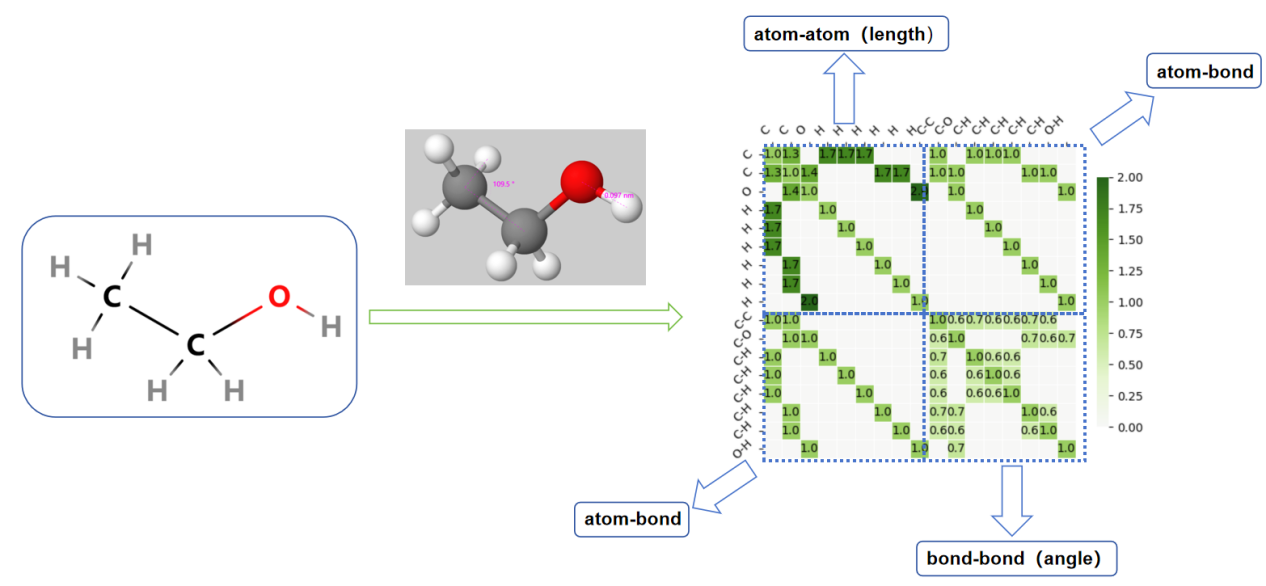

**Figure S1** Visualization of the three-dimensional spatial position matrix corresponding to the molecule of ethanol. Four matrices are used to represent atomic and bond relationships. Colors from light green to dark green indicate an increase in angle or distance.

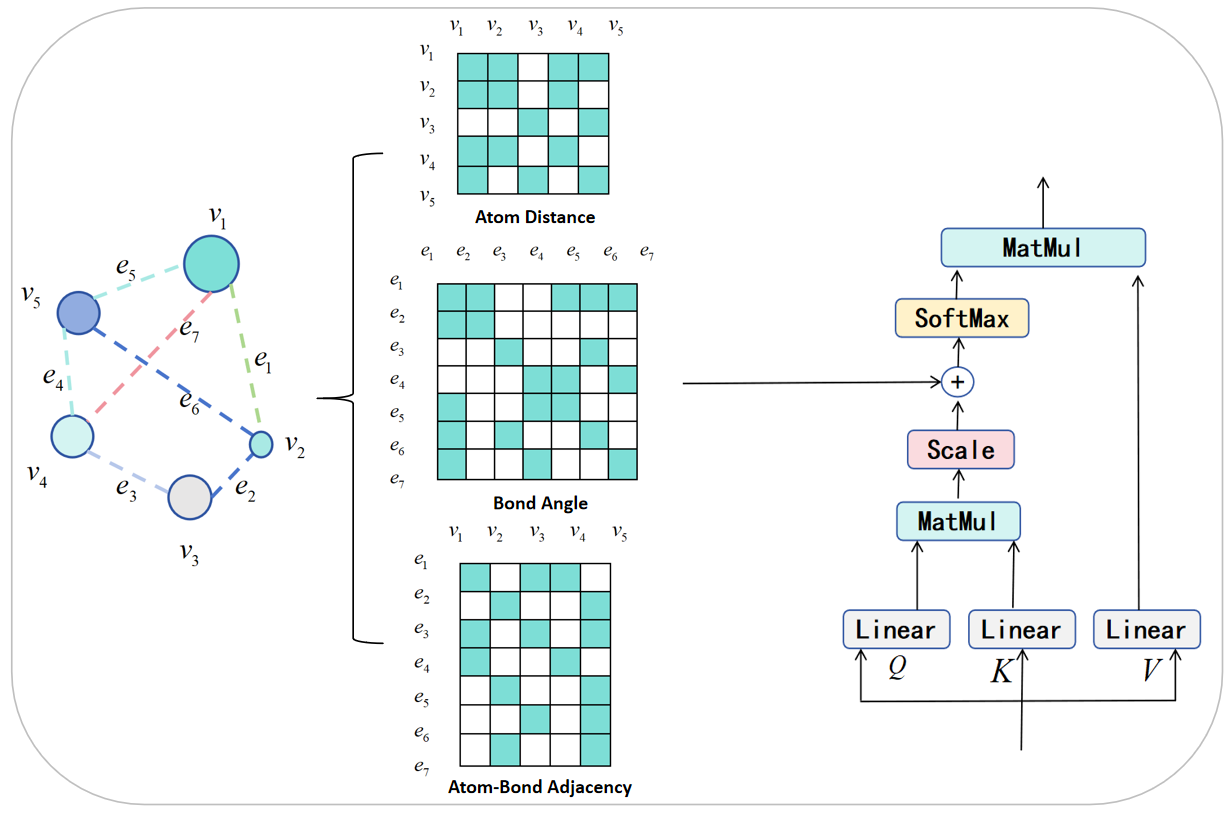

**Figure S2.** GEO-BERT [a] omits the Atom Distance and Bond Angle matrix, using a 0-1 mask matrix instead. When there is a bond between two atoms, the corresponding position is set to 1 on the Atom Distance matrix. Otherwise, set it to 0. When either end of the bond has the same atom, the corresponding position is set to 1 on the Bond Angle matrix. Otherwise, it is set to 0.

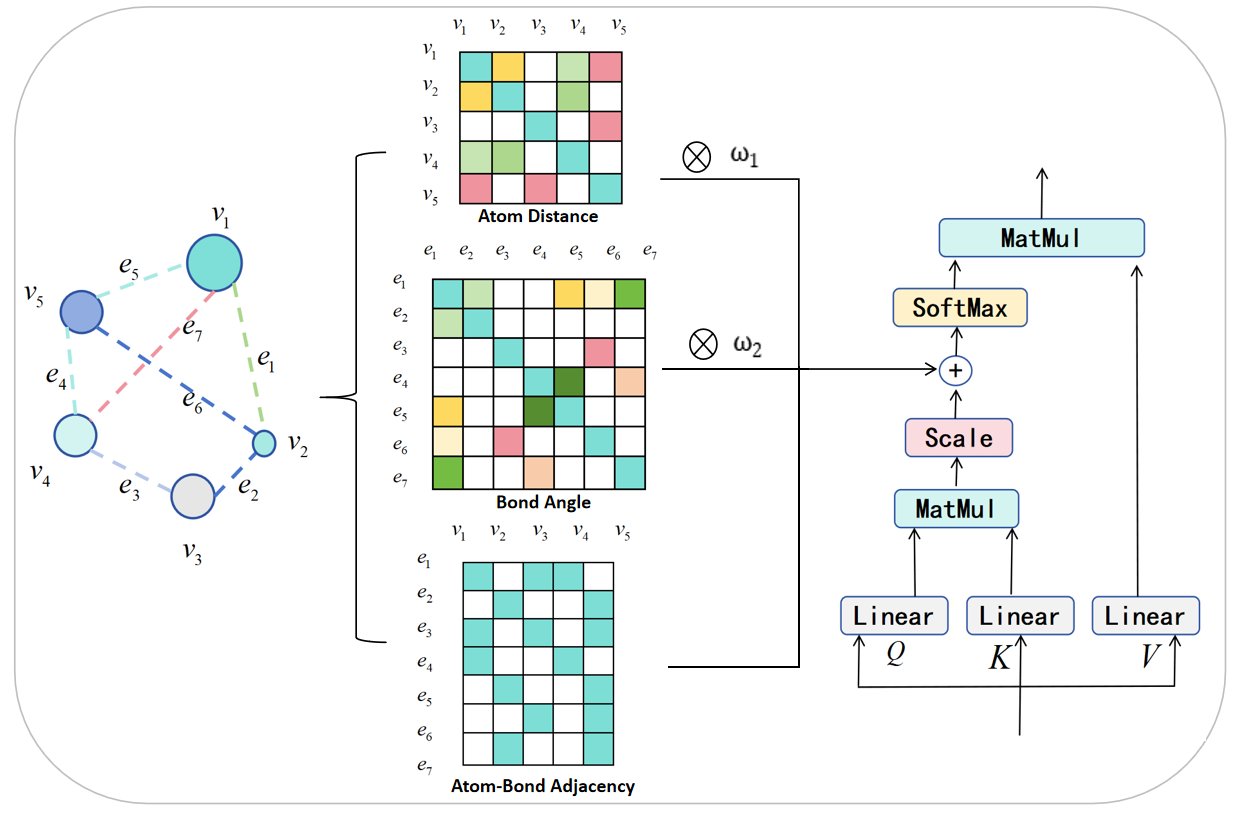

**Figure S3.** GEO-BERT [b] incorporates the atom distance matrix and bond angle matrix by first multiplying each of them by a learnable parameter and then adding them to the original attention matrix. These learnable parameters are shared across all the layers of GEO-BERT.

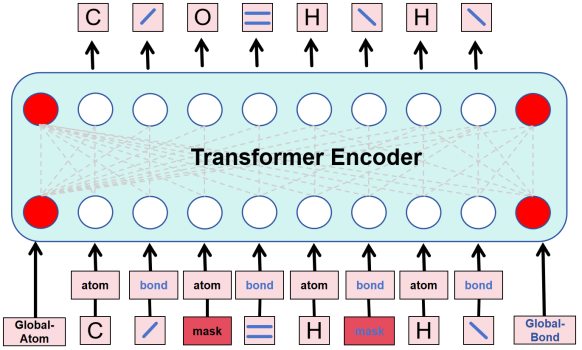

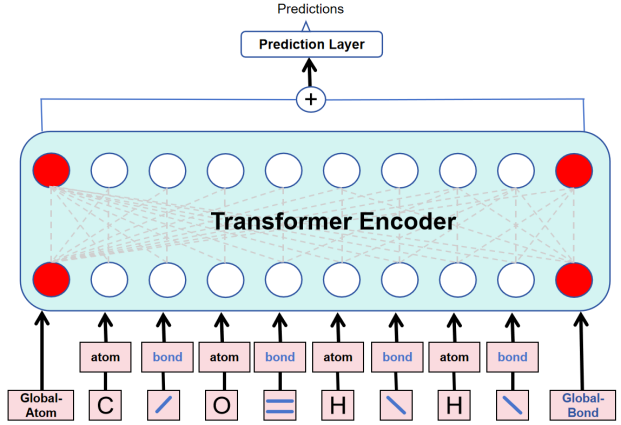

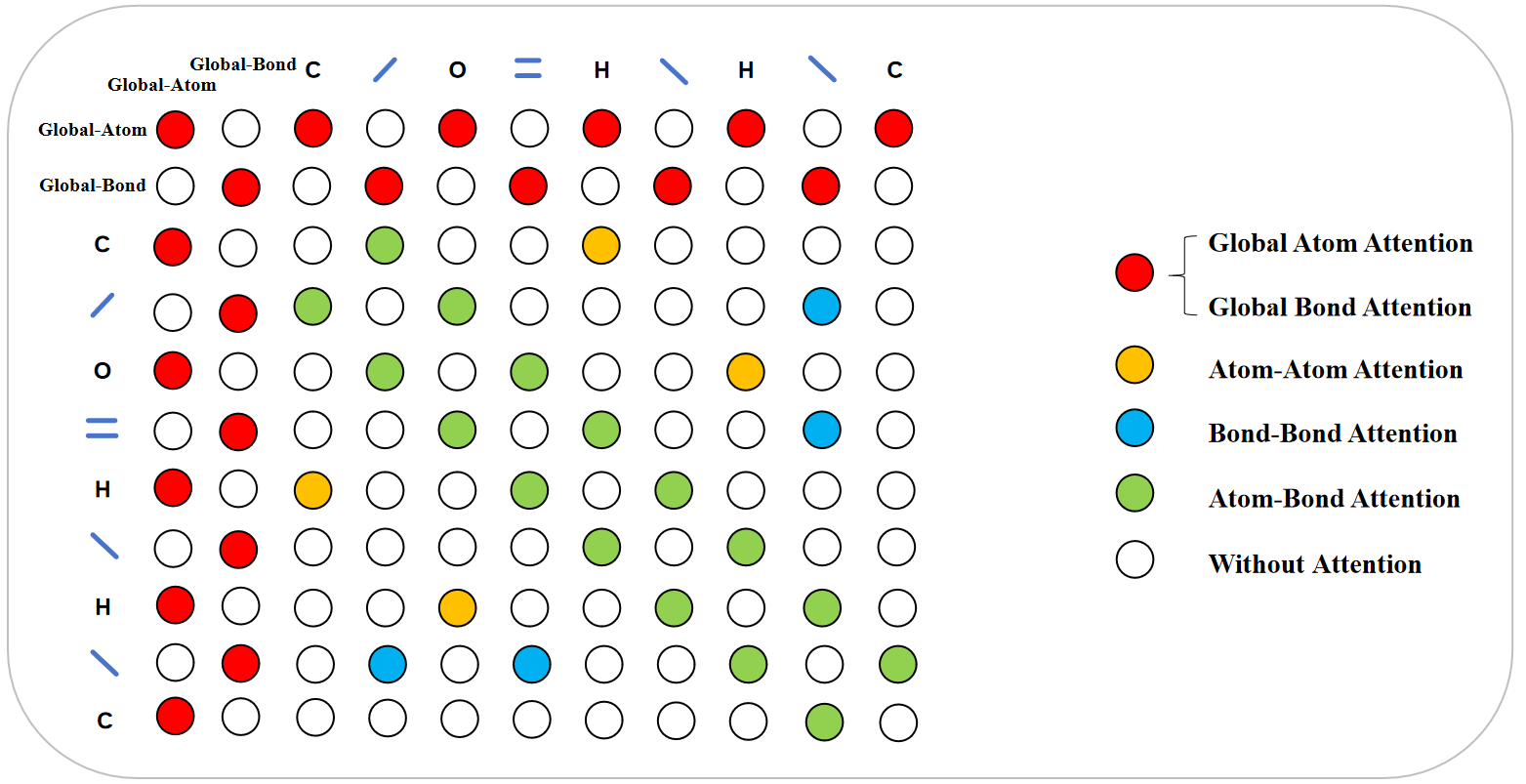

**Figure S4**. GEO-BERT [c] employs two special tokens (GLOBAL-ATOM and GLOBAL-BOND) to focus on the atoms and bonds in the molecule respectively, and subsequently concatenates the resultant vectors during the fine-tuning phase.

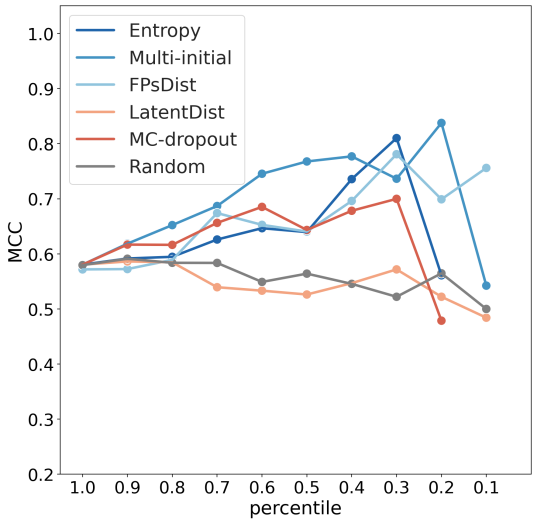

**Figure S5.** Uncertainty analysis of GEO-BERT(DYRK1A), based on other six uncertainty metrics and the “DYRK1A_test” dataset.

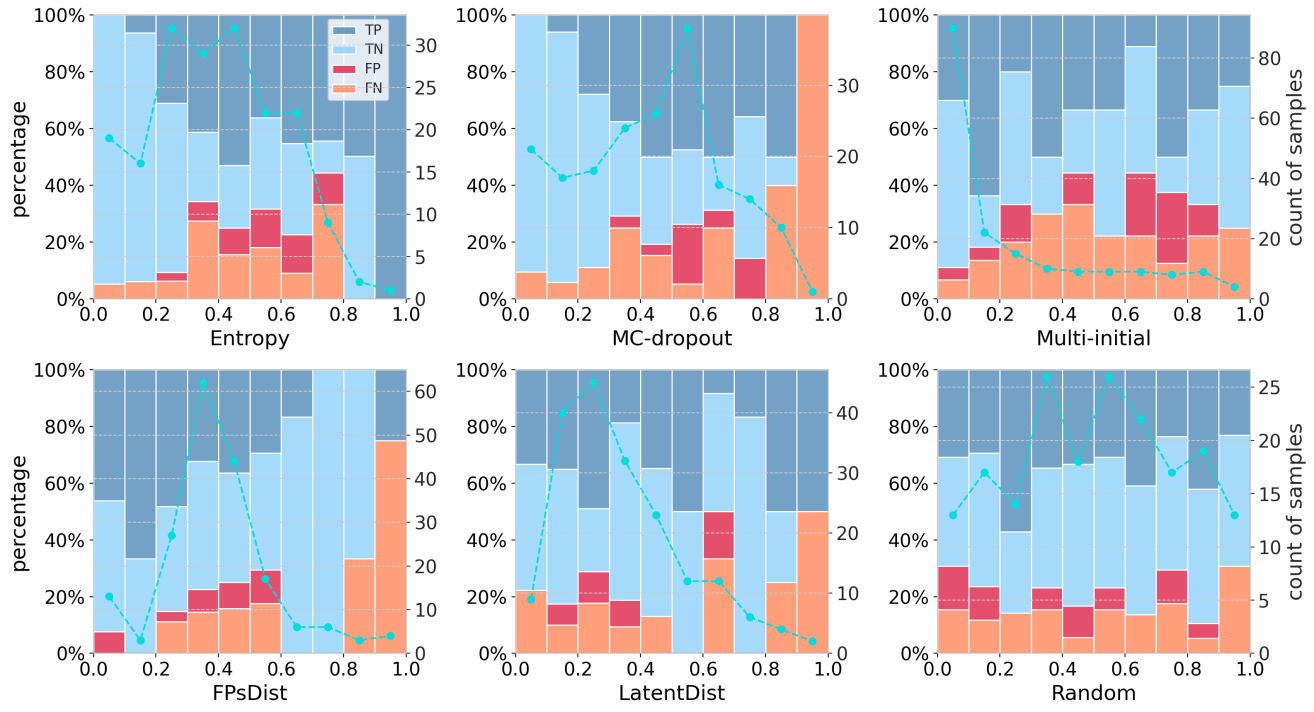

**Figure S6.** Distribution of various types of samples in the GEO-BERT model under six uncertainty indicators.

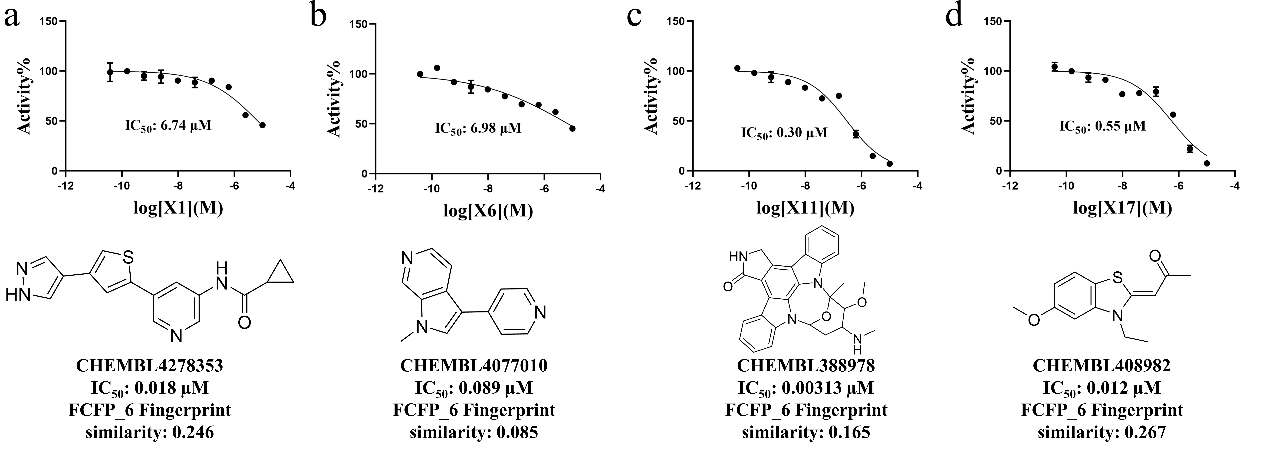

**Figure S7.** The inhibitory activity and structural novelty of compound **X1** (a), **X6** (b), **X11** (c) and **X17** (d), with Harmine as the reference compound (IC_50_: 21.24nM).

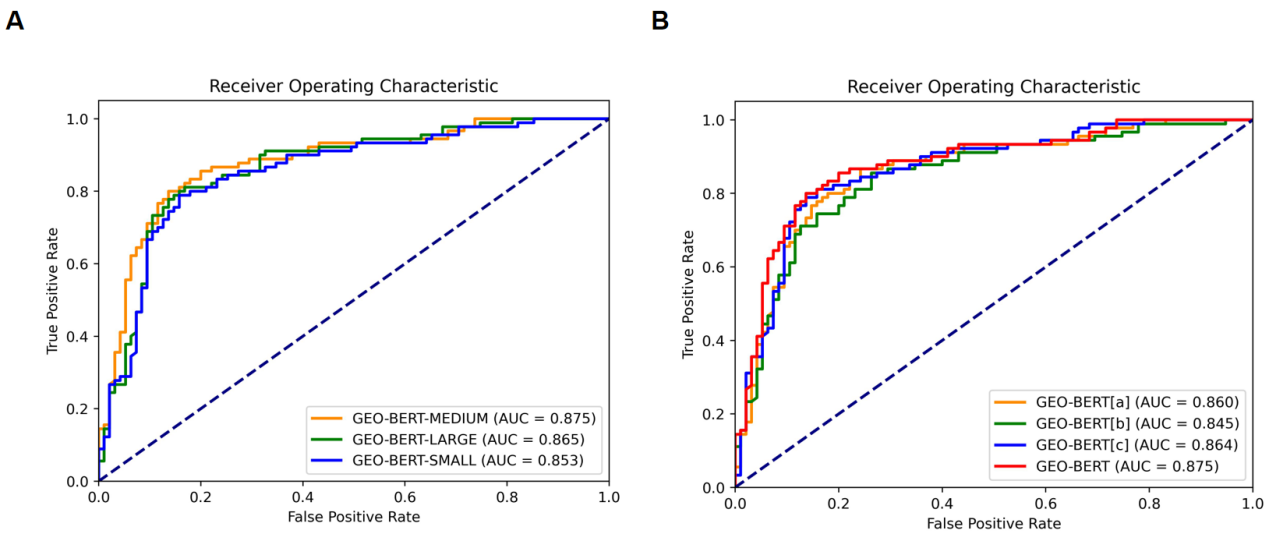

**Figure S8.** ROC curves of GEO-BERT on the DYRK1A dataset. (A) ROC curves of GEO-BERT with different scale sizes: GEO-BERT-[SMALL](AUC=0.853), GEO-BERT-[MEDIUM](AUC=0.875) and GEO-BERT-[LARGE](AUC=0.865). (B) ROC curves of various variants of GEO-BERT: GEO-BERT[a](AUC=0.860), GEO-BERT[b](AUC=0.845), GEO-BERT[c](AUC=0.864), and GEO-BERT(AUC=0.875).
